# Mesenchymal-Epithelial Transition Regulates Initiation of Pluripotency Exit before Gastrulation

**DOI:** 10.1101/655654

**Authors:** Sofiane Hamidi, Yukiko Nakaya, Hiroki Nagai, Cantas Alev, Takeya Kasukawa, Sapna Chhabra, Ruda Lee, Hitoshi Niwa, Aryeh Warmflash, Tatsuo Shibata, Guojun Sheng

## Abstract

The pluripotent epiblast gives rise to all tissues and organs in an adult body. Its differentiation starts at gastrulation when the epiblast generates mesoderm and endoderm germ layers through a process called epithelial-mesenchymal transition (EMT). Although gastrulation EMT coincides with loss of epiblast pluripotency, pluripotent cells in development and in vitro can adopt either mesenchymal or epithelial morphology. The relationship between epiblast’s cellular morphology and its pluripotency is not well understood. In this work, using chicken epiblast and mammalian pluripotency stem cell (PSC) models, we show that PSCs undergo a mesenchymal-epithelial transition (MET) prior to EMT-associated pluripotency loss. Epiblast MET and its subsequent EMT are two distinct processes. The former, a partial MET, is associated with reversible initiation of pluripotency exit; whereas the latter, a full EMT, is associated with complete and irreversible pluripotency loss. We provide evidence that integrin-mediated cell-matrix interaction is a key player in pluripotency exit regulation. We propose that epiblast partial MET is an evolutionarily conserved process among all amniotic vertebrates and its developmental function is to mediate planar symmetry-breaking within an epithelialized epiblast, taking place after epiblast MET but before gastrulation EMT.

## Introduction

A human embryo at late blastocyst stage of development contains three cell populations: the CDX2^+^ trophectoderm, GATA6^+^ primitive endoderm and POU5F1^+^ epiblast (1, 2). Among them, the epiblast is the only pluripotent population which will give rise to all cell lineages in an adult body. Implantation takes place soon afterwards and loss of pluripotency coincides with the onset of gastrulation when the epiblast initiates lineage differentiation by generating the three definitive germ layers (ectoderm, mesoderm and endoderm). Between pre-implantation and gastrulation (about one week in human development), the epiblast is considered to be pluripotent throughout. But due to ethical and technical limitations, morphogenesis of the epiblast during this period of human development is poorly understood. Different states of pluripotency maintenance in vitro have been hypothesized to correspond to unique sub-stages of pre-gastrulation epiblast morphogenesis, e.g., with the naïve and primed states representing the pre-implantation and post-implantation epiblast, respectively (3).

Pluripotency markers (e.g., NANOG and POU5F1) are expressed in both pre- and post-implantation epiblast and in their corresponding states captured in vitro in the mouse (4). The human epiblast is presumed to behave in a similar way, a concept partially supported by data from prolonged in vitro culture of human blastocyst and from histological analysis of rare, post-implantation stage human embryos (2, 5–7). However, differences between human and mouse development have long been noted, e.g., in topographic arrangement of the epiblast sheet with respect to the rest of the embryo in vivo (8) and in the properties of the embryonic stem cells (ESCs) and induced pluripotent stem cells (iPSCs) cultured in vitro (9). Furthermore, it is unclear whether epiblast’s intercellular organization plays any role in regulating its pluripotency in either mouse or human development. Except for germ cells, differentiation into any somatic cell lineage can be achieved from either the naïve or primed ESCs, suggesting that morphogenetic status of pre-gastrulation epiblast is not a key factor in its pluripotency maintenance in vitro. Yet, in all amniote species examined, including the birds and mammals, an epithelialized epiblast is a prerequisite for gastrulation to take place in vivo (8), suggesting that epiblast pluripotency status is causally correlated with its morphogenesis.

Epithelialization of mouse epiblast was shown to be dependent on cell-extra cellular matrix (ECM) interactions. Its polarization, for example, was reported to be regulated by Integrin-linked kinase activity, known to bridge β1/2/3-integrin cytoplasmic tail to the actin cytoskeleton (10), and by both β1-Integrin and Dystroglycan, two main types of transmembrane proteins that mediate epiblast-ECM interaction (11). Modulating cadherin-mediated epiblast cell-cell interactions, by either deleting E-cadherin or replacing E-cadherin with N-cadherin, also had a profound effect on epiblast cell-ECM affinity (12). Weakening of integrin-mediated cell-ECM interaction led to a reduction in E-cadherin mediated cell-cell adhesion strength (13), indicating that cell-ECM and cell-cell interactions, as well as their crosstalk, are critically involved in epiblast epithelialization (14). These lines of evidence suggest that polarization of the epiblast, including the establishment of apicobasal polarity and epiblast-ECM interaction and the modulation of epiblast adherens junction, is temporally and morphogenetically involved in epiblast pluripotency regulation.

To understand how epiblast morphology regulates its pluripotency, we first investigated whether its epithelialization, from late blastocyst to pre-gastrulation stage, could be viewed as a mesenchymal-epithelial Transition (MET) process, the reverse of Epithelial-Mesenchymal Transition (EMT) (15). Using a combination of in vivo (chicken epiblast) and in vitro (human iPSCs [hiPSCs], human ESCs [hESC] and mouse PSCs [mESCs and mEpiSCs]) models, we asked whether epiblast MET played a role in epiblast pluripotency maintenance and if so, how this could offer us insight into normal pre-gastrulation development of the human embryo. We present data showing that amniote epiblast goes through a partial MET process which is primarily characterized by the segregation of basal and lateral plasma membrane domains of the epiblast and by the deposition of epiblast basement membrane. This partial MET regulates the initiation of pluripotency exit through activation of the integrin-mediated signaling pathway.

## RESULTS

### Avian epiblast undergoes epithelialization and initiates pluripotency exit before the onset of gastrulation

Using the analogy of Waddington’s epigenetic landscape of lineage specification (16), the loss of pluripotency in amniote (birds and mammals) development is marked by gastrulation, a process in which the epiblast gives rise to the three definitive germ layers (the ectoderm, mesoderm and endoderm). We have previously shown that gastrulation in chick embryos, taking place mainly at Hamburger and Hamilton stages (HH) 3 and 4 (about 12-18 hours of development after egg-laying), requires an epithelial-mesenchymal transition (EMT) process to convert epithelial-shaped epiblast cells into mesenchymal-shaped mesendoderm precursors (17, 18). We and others have also reported that gastrulation EMT is associated with the loss of pluripotency markers in the epiblast at about stage HH5 when neural ectoderm fate is specified (19, 20). To investigate the dynamic relationship between epiblast EMT and its pluripotency loss, we carried out a transcriptomic analysis of pre- and peri-gastrulation stage chicken embryos (HH1-HH3; 0.5h-14h of development) (Fig.S1A). Area pellucida (intraembryonic) epiblast tissues, excluding the forming primitive streak, were collected and used for Affymetrix genechip-based transcriptome analysis (Materials and Methods). Differential expression analysis (Table S1) showed that 5529 (of 38535 in total) probe sets had statistically significant cross-stage variations (FDR<0.01), with 2376 of them exhibiting an increasing trend (Fig.S1B; 0001, 0011 and 0111 clusters) and 2543 a decreasing one (Fig.S1C; 1000, 1100 and 1110 clusters). Gene ontology analysis revealed a clear enrichment of biological processes associated with epithelial morphogenesis from stage HH2 onward (0001 cluster) (Table S2). Conversely, a large number of early HH1-specific genes (cluster 1000) were associated with RNA and nucleic acid binding, modification and metabolism (Table S2), likely reflecting their involvement in pluripotent maintenance at ovipositional stages as we have previously reported (21). This was supported by the analysis of candidate genes known to be involved in pluripotency regulation (e.g., NANOG, SOX3, FLF3, OTX2, TCF7L2, MYC, DNMT3 and LIN28) and epithelial morphogenesis (e.g., COL4 genes, SDC genes) (Fig.S1D; Table S1). Further confirmation came from re-analysis of our recently published developmental promoterome datasets (22). Although whole embryos, including both the area pellucida and area opaca, were used in that paper, a general decrease in promoter activities of pluripotency-related genes and increase in those of epithelial genes were observed prominently in the first day of chick development (HH1-HH7) (Fig. S2). Taken together, these data suggested that in addition to the well-known phase of pluripotency loss which is associated with gastrulation EMT and takes place at stage HH3-HH5, a decrease in pluripotency marker expression is seen much earlier from stage late HH1 and is possibly associated with an increase in epithelial-associated features of the epiblast.

### Avian epiblast is characterized by a partial mesenchymal-epithelial transition (MET) at late HH1 followed by a full EMT during gastrulation at HH3

To better characterize this epithelialization process, we analyzed stage HH1 embryos with markers associated with epithelial polarity (Fig.1). We compared embryos from freshly-laid eggs, at Eyal-Giladi and Kochav (23, 24) (EGK-) X/XI sub-stages of HH1 (referred to as early HH1 in this work), with embryos from eggs incubated for six hours, at EGK-XIII/XIV substages of HH1 (referred to as late HH1). At late HH1, epiblast polarity was clearly established, as evidenced by the presence of a continuous layer of basement membrane (Laminin and Agrin; Fig.1A), presence of apical membrane domain (aPKC; Fig.1G,H) and apical tight junctions (ZO-1; Fig.1E,F), segregation of basal and lateral membrane domains marked by the lateral localization of E-cad (Fig.1B) and basal enrichment of beta1 Integrin and Dystroglycan (Fig.1C,D), and enrichment of Golgi apparatus (GM130; Fig.1I,J) and acetylated tubulin (Fig.1K,L) in the cytoplasm apical to the nucleus. Interestingly, all these markers were also expressed at early HH1, but at lower levels and/or in a much less organized fashion (Fig.1; 0 hr). For example, in early HH1 embryos, Laminin and Agrin were expressed spottily under a small percentage of epiblast cells (Fig.1A) and E-cad marked both lateral and basal regions of an epiblast cell (Fig.1B). Both ZO-1 and aPKC exhibited apically-compartmentalized yet uneven distribution at early HH1 (Fig.1E,G). Taken together, these data suggested that at early HH1 (egg-laying) chick epiblast cells exhibit weak epithelial features and they become fully epithelialized in about six hours of development when the embryo reaches late HH1 stage (Fig.S3), approximately six hours before the onset of gastrulation at HH3. This transition from a weakly polarized organization to a fully epithelial organization can be viewed as epithelioid to epithelial transition (referred to as a partial mesenchymal-epithelial transition in this work; partial MET). We then used Laminin expression as an indicator of the extent of this partial MET and NANOG protein expression as an indicator of epiblast pluripotency and investigated their relative and dynamic changes from HH1 to HH3 (Fig.S4A-D). Before full epithelialization, a small increase in NANOG levels was observed from early HH1 to late HH1 (Fig.S4A,B). This likely corresponded to the final step of epiblast-hypoblast sorting through polyingression which was reported in the chick (25–27) and which in the mouse was associated with a loss of GATA6 expression and an increase of NANOG expression (28). After full epithelization, NANOG levels in epiblast cells decreased steadily from late HH1 (Fig.S4B) to late HH2 (Fig.S4C) and late HH3 (Fig.S4D). As shown previously (19), after gastrulation EMT, mesendoderm cells lost their pluripotency marker expression completely (Fig.S4D).

**Figure 1:**
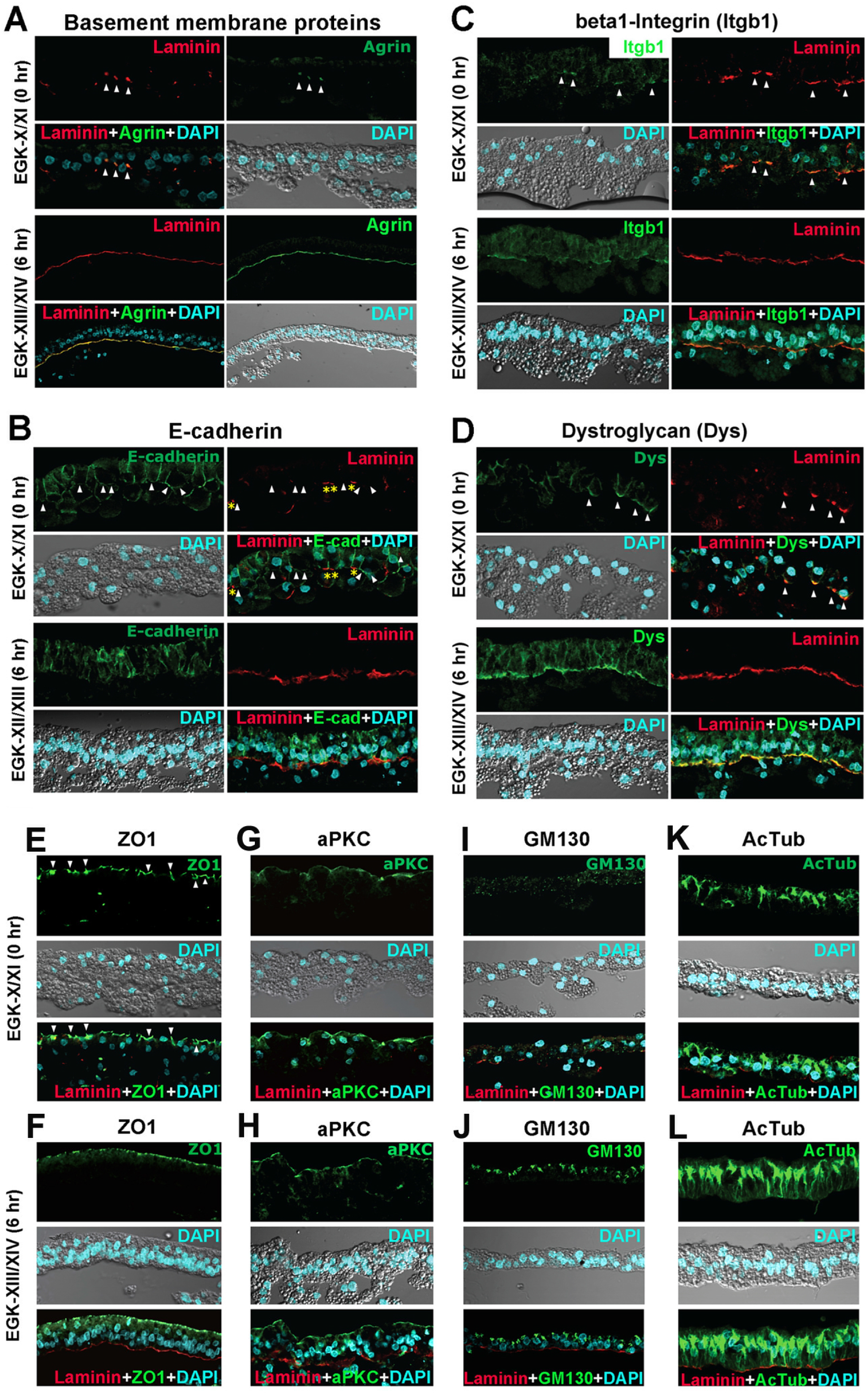
Immunofluorescence staining of stage HH1 chicken epiblast with epithelial markers. (A) Laminin (red), Agrin (green) and DAPI (cyan) staining with unincubated (EGK-X/XI) and 6-hour incubated (EGK-XIII/XIV) embryos. AT EGK-X/XI, basement membrane is absent or very immature (Arrowheads). At EGK-XIII/XIV, basement membrane is mature. (B) Laminin (red), E-cadherin (green) and DAPI (cyan) staining with unincubated (EGK-X/XI) and 6-hour incubated (EGK-XIII/XIV) embryos. AT EGK-X/XI, distinction between basal and lateral membrane domains is not clear and E-cadherin is expressed in both domains (arrowheads). In epiblast cells with basement membrane deposition, basal E-cadherin localization is suppressed (asterisks). Epiblast cell height is uneven. At EGK-XIII/XIV, most of epiblast cells have basement membrane underneath them and have laterally restricted E-cadherin expression. (C) Laminin (red), beta1 Integrin (green) and DAPI (cyan) staining with unincubated (EGK-X/XI) and 6-hour incubated (EGK-XIII/XIV) embryos. (D) Laminin (red), Dystroglycan (green) and DAPI (cyan) staining with unincubated (EGK-X/XI) and 6-hour incubated (EGK-XIII/XIV) embryos. (E, G, I, K): Un-incubated embryo (EGK-X/XI) stained for tight junction marker ZO-1 (E), apical marker aPKC (G), Golgi marker GM130 (I) and acetylated tubulin (K). (F, H, J, L): Embryos incubated for 6 hours (EGK-XIII/XIV) stained for tight junction marker ZO-1 (F), apical marker aPKC (H), Golgi marker GM130 (J) and acetylated tubulin (L). All markers were co-stained for Laminin (red) and DAPI (cyan).

### Undifferentiated hiPSCs exhibit a size-dependent shift in macroscopic patterns of pluripotency gene expression

Similar to the avian epiblast, mammalian pluripotent cells appear also to undergo an epithelialization process in their morphogenesis, i.e., from non-epithelial epiblast precursors in the inner cell mass to an epithelial epiblast surrounding a proamniotic cavity before gastrulation (29). We hypothesized that such a partial MET, similar to the case in the chick, is critical for mammalian pluripotency regulation, possibly marking a so far undescribed initiation checkpoint of pluripotency exit. To test this hypothesis, we used cultured human induced pluripotent stem cells (hiPSCs; 201B7 line) (30) as a surrogate for the human epiblast tissue. HiPSCs can be kept in a pluripotent state under maintenance conditions and can give rise to cell lineages in all three germ layers upon receiving differentiation cues. Single-cell level heterogeneity in pluripotency marker expression has been reported for hiPSCs cultured under maintenance conditions (31–33).

We first asked whether a similar heterogeneity could be observed at the macroscopic level. To reveal dynamic changes in gene expression, we used the RNA in situ hybridization method and analyzed expression patterns of key pluripotency regulatory genes POU5f1, NANOG, LEF1 and OTX2 in hiPSCs (Materials and Methods; Fig.2A). For all four genes, we observed two types of hiPSC colonies (Fig.2A), one with a ubiquitous and centrally-high expression [referred to as center (+) colonies in this work] (Fig.2A, top panels; Fig.2C) and the other with a pericentrally-high, but centrally-low or –negative expression [referred to as center (-) colonies in this work] (Fig.2A, bottom panels; Fig.2C). Statistical analysis revealed that center (+) colonies were generally smaller (with a colony radius of 304.6 +/-12.3 μm and n=53 for POU5F1) than the center (-) ones (with a colony radius of 553.7 +/-27.6 μm and n=30 for POU5F1) (Fig.2A,B). Similarly, a strong size-pattern correlation was observed for the other three genes (Fig.2B). To understand this dynamic shift in more detail, we divided each POU5F1-stained colony into sub-territories as schematized in Fig.2C and analyzed their correlation with the colony size (Fig.2D). The center (-) territory was strongly correlated with the overall colony size (Fig.2D green) (Pearson correlation coefficient r= 0.88; two-tailed t-test p< 0.001). A clear correlation (r= 0.72, *p< 0.001*) was also seen between the colony size and the width of POU5F1-positive territories [center (+) region in small colonies and pericenter (+) region in big colonies as shown in Fig.2D], with an initial positive association (up to 450 μm in colony size) and a gradual shift to a steady state width of approximately 150 μm (Fig.2D blue). On the contrary, no correlation between the colony size and the edge territory was observed (r=0.22, *p=ns*) (Fig.2C; Fig.2D purple). This overall dichotomy in expression patterns was further supported by POU5F1 and NANOG immunofluorescence analysis (Fig.2E). Taken together, these data suggested that hiPSCs cultured under pluripotency maintenance conditions undergo a macroscopically predictable change in pluripotency gene expression during their expansion from a few cells to a colony of hundreds of cells before passaging.

**Figure 2:**
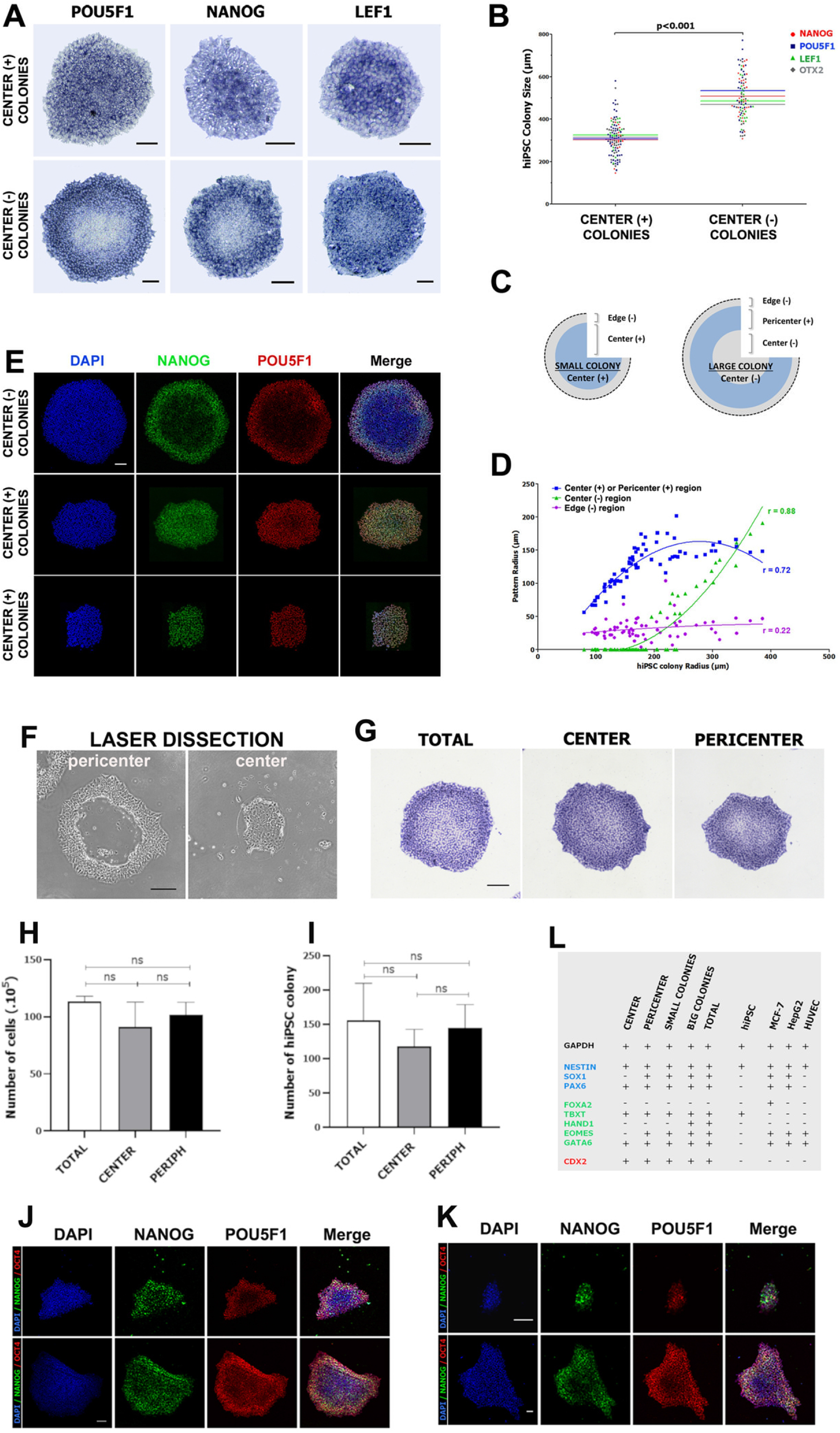
Undifferentiated hiPSCs exhibit dynamic patterning of pluripotency gene expression without pluripotency loss. (A) RNA in situ hybridization analysis of POU5F1, NANOG, and LEF1 genes in hiPSC colonies at day 6 of culture, revealing two types hiPSC colony organization: central (+) or central (-). Details on each type shown in panel (C). (B) Correlation of colony size (diameter in μm) and colony type [center (-) or center (+)] for each gene (NANOG, POU5F1, LEF1 and OTX2). (C) Schematic organization of two colony types. Small colonies (^~^300 μm in diameter) are center (+) and large colonies (^~^500 μm in diameter) are center (-). In each case, there is an “edge” sub-territory which has low RNA expression associated with flattened cell morphology. This edge (-) sub-territory defines the boundary of colony and is not studied in detail in this work. (D) Correlation analysis between the width (pattern radius) of those three sub-territories and hiPSC colony radius. “r”: Pearson correlation coefficient. Lines correspond to nonlinear polynomial of order 2 (quadratic) regression curve based on the plotted points. (E) Confocal images of NANOG and POU5F1 protein expression in hiPSC colonies of increasing sizes (bottom to top). Scale bar: 100 μm (for all panels). (F) Laser-assisted microdissection of center and pericenter cells in patterned large hiPSC colonies. (G) Pou5F1 RNA After re-culture (by passaging) of microdissected cells, both center-derived and pericenter-derived cells were able to reform size-dependent colony patterning, with no clear difference from each other and from whole colony re-culture. (H, I) Statistical analysis of cell number and colony number after re-culture. No significant difference, either in number of post-culture cells (H) or in number of colonies formed (I), could be seen between total-, center- and pericenter-derived cells. (J, K) After laser-microdissection, remaining cells in the colony, either as center cells (K) or as half a colony (J), could continue colony growth and reform a patterned colony. (L) Without re-culture, differentiation potentials of the center cells were favored towards the ectoderm lineage. Micro-dissected hiPSCs were collected, aggregated and induced to differentiate (see Materials and Methods), followed by expression analysis of ectoderm-, mesoderm- and endoderm-lineage specific markers. Center-derived hiPSCs had a reduced ability to form mesoderm lineages.

### Reduction in pluripotency gene expression in hiPSC colonies is a reversible process with no loss of pluripotency after re-culture

To test whether reduced pluripotency gene expression in center (-) colonies was indicative of an irreversible pluripotent loss, we isolated center and pericenter cells through laser microdissection (Fig.2F) (materials and methods). Isolated cells were treated with ROCK inhibitor Y-27632 for 2h before dissociation and re-culture in normal maintenance media. Both center-derived and pericenter-derived hiPSCs were capable of growth and re-formation of the originally patterned colonies (Fig.2G) with no significant difference in either the total cell number (Fig.2H) or colony number (Fig.2I) after re-culture. When large colonies were micro-dissected to remove pericenter cells, leaving only center (-) cells (Fig.2K, top), such colonies were able to reform large and patterned colonies after continuation of the culture without passaging (Fig.2K, bottom). Similarly, bisected colonies (with half of the colony removed) were also able to re-establish patterning after continuation of the culture (Fig.2J). Those data showed that hiPSCs located in the colony center, although having much reduced levels of pluripotency gene expression, still retained their pluripotency and that such pluripotency could be manifested after dissociation or disruption of original colony organization. However, center and pericenter cells did exhibit biased differentiation capability (Fig.2L). When we collected micro-dissected center and pericenter cells for direct differentiation (Materials and Methods), pericenter cells had full capability (compared with non-dissected hiPSC colonies) to differentiate into the three germ layers, whereas center cells had reduced capability to differentiate into mesoderm fate (Fig.2L).

### The decrease in hiPSC pluripotency is correlated with an increase in its epithelialness

Full pluripotency loss takes place during gastrulation EMT. Our data on the chicken epiblast suggested that an important landmark of pluripotency exit is the transition from a non-epithelial epiblast to an epithelial epiblast (i.e., epiblast MET rather than EMT). We asked whether this macroscopic shift in pluripotency gene expression in hiPSC colonies described above was correlated with changes in their epithelial status. Close observation indicated that in small colonies, cells in the center were loosely packed (Fig.3A,C). In large-sized colonies, only colony-pericenter cells showed similar loose packing, whereas colony-center cells were arranged tightly (Fig.3B,D). This was supported by histological sections of hiPSC colonies cultured on polycarbonate membrane (Fig.3E, top and middle panels). A unilaminar (single-cell layered) organization of hiPSCs, regardless of colony size, was also evident from histology and confocal analyses (Fig.3E,H-K). Flattened cells at the colony edge (Fig.3A-E) were POU5F1 RNA-low cells (labelled as edge (-) in Fig.2C). Those cells defined colony boundary, were involved in colony size expansion, and were not studied further in this work. Intercellular space seen in loosely-packed cells (center cells in small colonies and pericenter cells in large colonies) (Fig.3A-E) was restricted to the basal side because little intercellular space was observed at the apical side of either tightly-packed cells in the center (Fig.3F, left) or loosely-packed cells in the pericenter (Fig.3F, right). Similar to the temporal progression of epithelial maturation described in the avian epiblast (Fig.1), a spatial progression of epithelial maturation was seen in hiPSCs (Fig.3G-K). Both ZO-1 (tight junction marker) (Fig.3G,H) and E-cad (adherens junction marker) (Fig.3G,I) were expressed in both center (tightly packed) and pericenter (loosely packed) cells. However, ZO-1 and E-cad were diffusely localized in pericenter cells (Fig.3H,I), characteristic of an immature epithelium (34, 35); whereas in center cells, E-cad (Fig.3I) was localized to the lateral membrane and ZO-1 (Fig.3H) to the apical junctions, as expected for a mature epithelial organization. The full epithelial nature of center cells was further supported by deposition of basement membrane proteins (HSPG2 in Fig.3J; LAMA1 in Fig.3K) under colony center cells, but not pericenter cells. Interestingly, apical membrane marker aPKC did not exhibit prominent difference in either expression level or localization between center and pericenter cells (Fig.3L), suggesting that the pericenter cells, although not fully epithelial, exhibit partial epithelial characteristics and that these cells will progressively mature into a full epithelium as the colony expands. Collectively, these data showed that hiPSCs under pluripotency maintenance culture conditions undergo a partial MET that is correlated with a reduction in their pluripotent gene expression. Taking both chick epiblast and hiPSC data into consideration, we called this phenomenon MET-associated initiation of pluripotency exit in order to distinguish it from the loss of pluripotency associated with gastrulation EMT.

**Figure 3:**
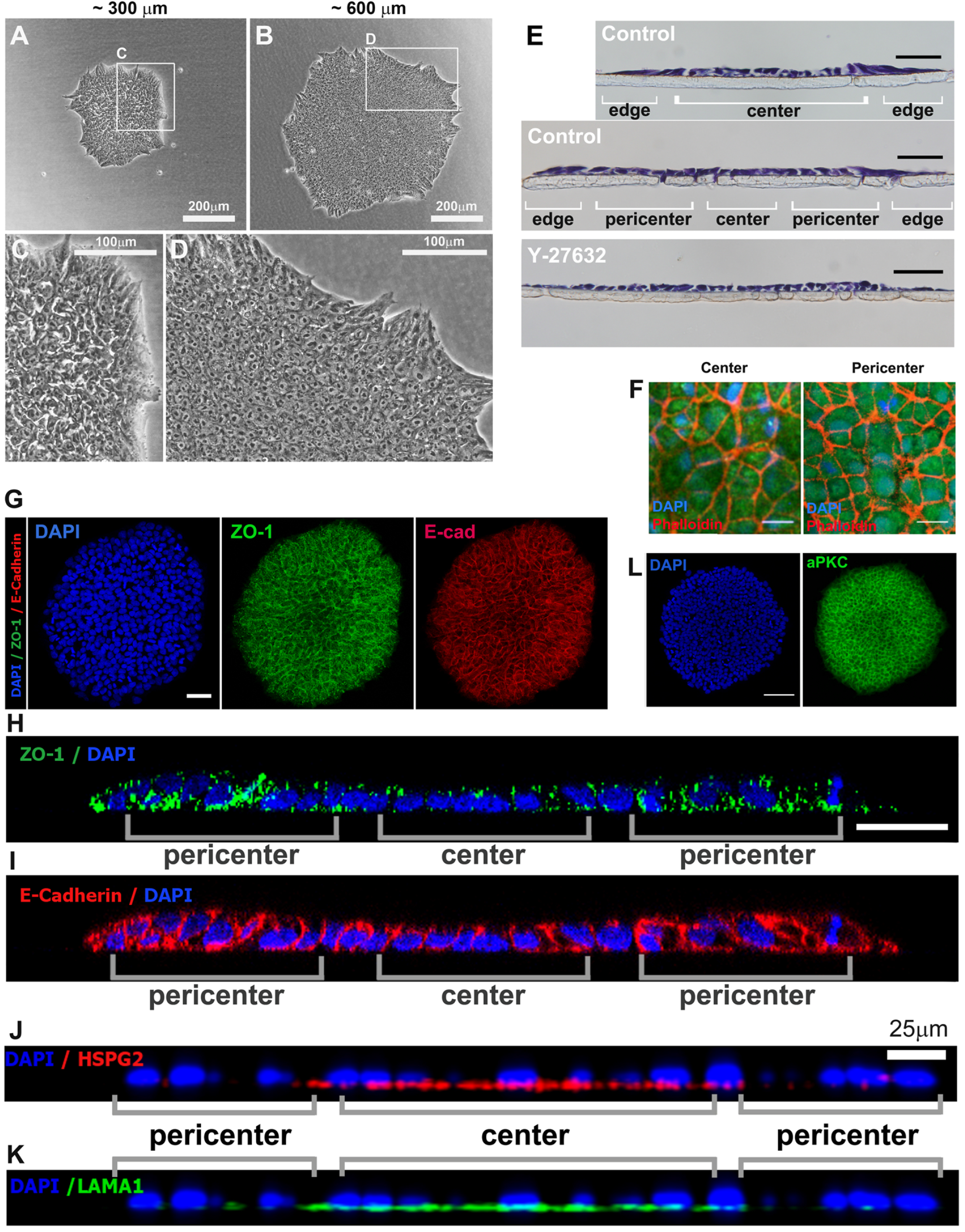
Epithelial status of hiPSC colonies. (A-D) Bright-field views of small (A; magnified view in C) and large (B; magnified view in D) hiPSC colonies, showing loose intercellular organization in small colonies (A, C) and tight intercellular organization in large ones (B,D). Scale bar: 200 μm in A and B; 100 μm in C and D. (E) Histological section of hiPSCs grown in polycarbonate membrane, supporting the observation in A-D. Top panel: small colony; middle: large colony; bottom: colony treated with ROCK inhibitor Y-27632. Scale bar: 40 μm. (F) Confocal images of the apical side of a large colony, showing its center (left panel) and pericenter (right panel) areas. In both areas, hiPSCs do not show any intercellular gap between cells. (G-K) Representative immunofluorescence images of hiPSC colony stained with tight junction marker ZO-1 (colony view in G, z-section view in H), adherens junction marker E-cadherin (colony view in G, z-section view in I), basement membrane marker HSPG2 (z-section view in J) and LAMA1 (z-section view in K), and apical marker aPKC (colony view in L). aPKC staining is uniform in the colony (L). ZO-1 and E-cad show a stronger, but less localized expression in colony pericenter. In colony center, both ZO-1 and E-cad exhibit epithelial-like localized organization, with ZO-1 at the apical junction and E-cad at the lateral membrane. HSPG and Laminin deposition is prominent only in colony center.

### Perturbation of MET alters initiation of pluripotency exit

To test whether the association between MET and pluripotency is causal, we used the Rho-kinase inhibitor Y-27632 to perturb the MET process. We have previously shown that the activity of small GTPase RhoA is essential for the maintenance of epithelial status of the epiblast during gastrulation EMT (17). Y-27632, a potent inhibitor of Rho-associated protein kinases and of RhoA activity, was reported to increase the survival of dissociated hiPSCs without significantly altering their pluripotency status (36). After 48 hours of Y-27632 treatment (20μM; from day 4 to day 6 of culture), cells in the center of hiPSC colonies adopted a loosely-packed morphology (Fig.3E, bottom panel) compared to the control (Fig.3E, middle panel), suggesting that the process of partial MET in the colony center was inhibited. Statistical analysis revealed that Y-27632 also caused a reduction in hiPSC’s height (Fig.4A; black control r=0.82 p<0.001, blue Y27632 r=0.50 p<0.001) and height heterogeneity (measured by max/min cell height ratio; Fig.4B; black control r=0.81 p<0.001, blue Y27632 r=0.20 p=ns). We then asked whether Y27632 could affect pluripotency exit as described above. After either 24 hours (day 5 to 6) or 48 hours (day 4 to 6) treatment with Y-27632, iPSC colonies were analyzed for POU5F1 expression. Two types of POU5F1-expressing colonies were observed in both cases (Fig.4C), similar to the control. A dramatic increase in colony size, however, was observed after both 24 hour and 48 hour treatment and in both colony types (Fig.4D for center-positive colonies and Fig.4E for center-negative colonies), consistent with the reduction in cell-cell compactness (Fig.3E) and cell height (Fig.4A,B). Moreover, in the 24 hour treatment group, the relative ratio of center-negative and pericenter-positive territories in large colonies was changed (Fig.4C,F), resulting in an increase in the POU5F1-positive pericenter territory at the expense of POU5F1-negative one (Fig.4C,F; green: 24 hour-treated; blue: control). This increase in the POU5F1-positive pericenter territory was correlated with the increase in the overall colony size (r=0.72 p<0.001) (Fig.4F). These observations were further supported by the analysis using POU5F1 and NANOG antibodies (Fig.4G). In the 48 hour treatment group, the size of POU5F1-positive territory became variable and lost any correlation with the colony size (r=0.21, p<0.05) (Fig.4C,F), although a mild increase throughout the colony in POU5F1 expression levels was observed for this group (Fig.4C). To test whether such perturbation of patterning was specific to Y-27632, we cultured hiPSCs for six days (the entire duration of culture) under hypoxia (5% O_2_), which had been reported to promote pluripotency maintenance in ESCs (37, 38). Hypoxia caused a small increase in overall colony size (Fig.4C-E), but without altering the patterning of the colony (Fig.4F; compare red and blue lines; r=0.56 and r= 0.61 respectively with p>0.001 in both conditions). Taken together, these data suggested that Y-27632 treatment disrupted the partial MET in hiPSC colonies and resulted in a delay in MET-induced pluripotency exit and a mild increase in pluripotency marker expression.

**Figure 4:**
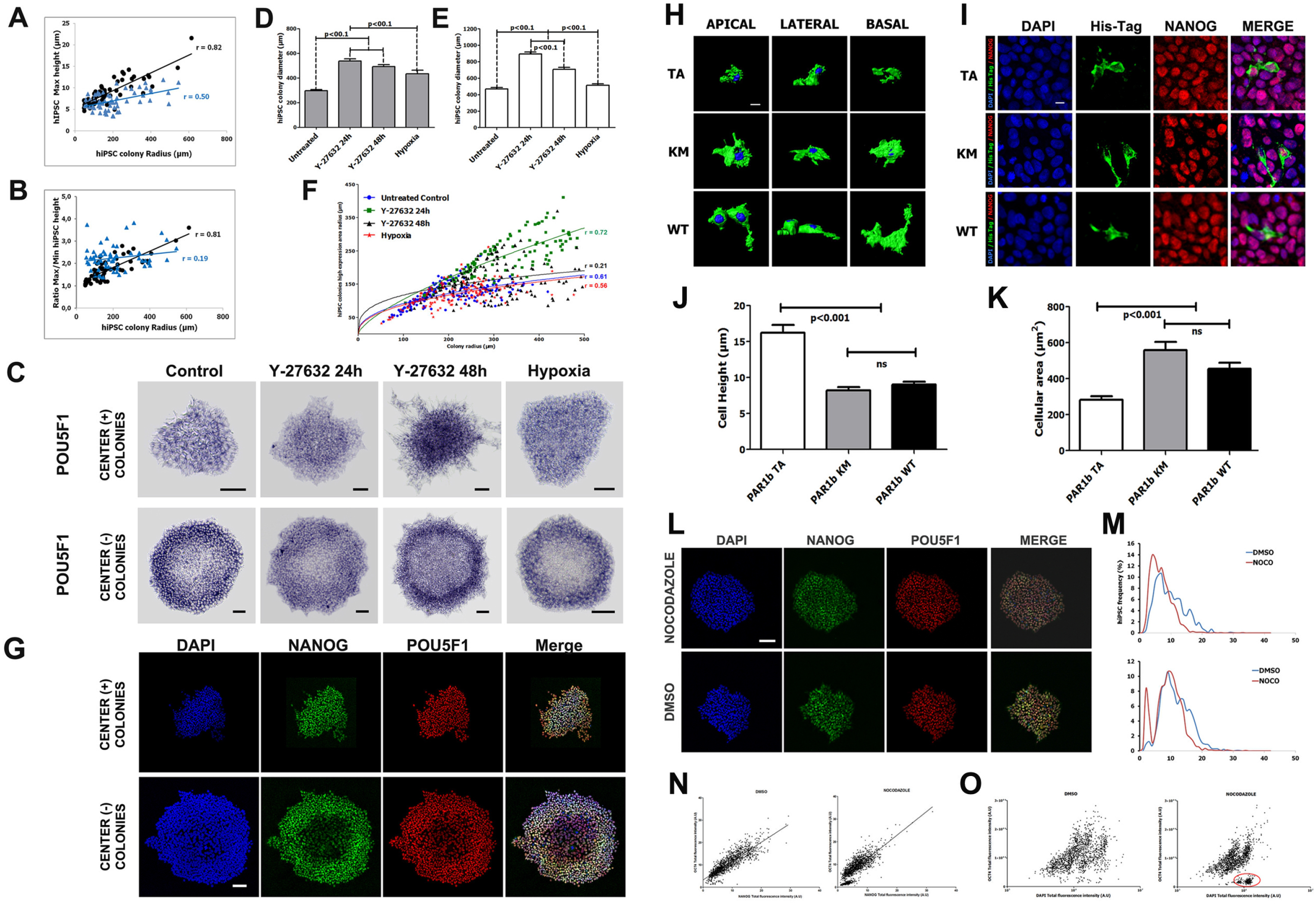
Perturbation of MET alters pluripotency gene expression and cell height modulation does not induce pluripotency ext. (A and B) Correlation plot between the cellular height and the colony radius in each hiPSC colony. “r”: Pearson correlation coefficient. Lines correspond to the linear regression based on the plotted point. Black: control colonies; Blue: Y-27632 treated colonies. (A) correlation of colony radius with maximum cell height, (B) correlation of colony radius with the ratio between maximum/minimum cell height. Y-27632 reduces hiPSC max height and completely disrupts intra-colony height ratio organization and its correlation with colony size. (C-F) RNA in situ hybridization analysis of POU5F1 gene in hiPSC colonies at day 6 of culture in control, Y-27632 treated (24h and 48h) and hypoxia-treated (5% O2 for six days) conditions. (C) Representative images of POU5F1 expression after treatment. (D-E) Analysis of central (+) (D) and center (-) (E) hiPSC colony diameter in these culture conditions. (F) Correlation analysis between the colony radius and the width of pluripotency gene expressing areas [center (+) or pericenter (+) sub-territories]. Lines correspond to a logarithmic nonlinear regression curve based on the plotted points. “r”: Pearson correlation coefficient. (G) Confocal images of POU5F1 and NANOG protein expression in hiPSC colonies treated with Y-27632 for 24h. Scale bar: 100μm. (H-K) Analysis of hiPSC morphological changes 48 hours after transfection of expression vectors for one of PAR1b protein mutant forms: PAR1b-TA, -KM and -WT (see Material and Methods). (H) Representative images of apical (left), lateral (middle) or basal (right) projected 3D reconstruction of hiPSC, showing that PAR1b-TA significantly increased hiPSC height and decreased apically or basally projected cell areas (quantification in J and K, respectively). Scale bar: 5μm. (I) Co-staining of PAR1b (His-tag) and NANOG, showing that NANOG expression is not altered by PAR1b over-expression. Scale bar: 5μm. (L-O) NANOG and POU5F1 expression after nocodazole treatment. (L) Representative confocal images of NANOG and POU5F1 protein expression after DMSO or nocodazole treatment. (M) Distribution analysis of NANOG (top panel) and POU5F1 (bottom panel) expression levels (in fluorescence A.U. here and in following graphs) for DMSO (blue) and nocodazole (red) treated cells, showing a modest decrease in fluorescence intensity after nocodazole treatment. (N, O) Correlation plots between POU5F1 and NANOG (N), and between POU5F1 and DAPI (O) expression levels. Line slope corresponds to the linear regression for those cells. Red circle in (O) highlights the DAPI high POU5F1 low population which is under cell-cycle arrest.

### Epithelial cell height is not a significant indicator of pluripotency exit

After epithelization but before gastrulation, the epiblast in both chicken and human embryos undergoes further morphogenetic changes, among which is an increase in cell height. During gastrulation EMT (pluripotency loss), epiblast cells are typically of columnar epithelial morphology. In this study, we observed heterogeneity in hiPSC height (Fig.4A; black control), although a simple correlation between cell height and pluripotency level could not be established. To test whether cell height plays an instructive role in pluripotency exit, we used a constitutively active version of human PAR1b (hPAR1b-TA; Materials and Methods) to regulate hiPSC height. PAR1 is a serine-threonine kinase important for cellular polarity establishment and maintenance. hPAR1b-TA (T595A), a non-phosphorylatable version of hPAR1b which renders it “constitutively active”, was shown to promote basolateral membrane domain at the expense of apical domain and to increase the cell height of MDCK monolayer (39). As expected, expression of hPAR1b-TA in hiPSCs (analyzed 48 hours after transfection) led to a dramatic increase in cell height (Fig.4H, lateral view; Fig.4J, quantification), as well as to a decrease in projected cell surface area (Fig.4A; apical and basal views; Fig.4K, quantification). However, co-immunofluoresence analysis of hPAR1b-TA (revealed by antibody against His-tag) and NANOG showed that no significant change in pluripotency level could be associated with increased cell height (Fig.4I). Neither wild type hPAR1b (hPAR1b-WT) nor a kinase-dead mutant of hPAR1b (hPAR1b-KM) had a clear effect on either cell height (Fig.4H,J,K) or pluripotency marker level (Fig.4I). These data, together with our observation that in both chick epiblast (Fig.1) and hiPSC colonies (Fig.3) the apical domain marker aPKC was already correctly localized at the immature epithelial stage, suggested that the segregation of apical and basolateral membrane domains was not a key factor in this partial MET or pluripotency exit.

### Integrin-mediated compartmentalization of lateral and basal domains regulates pluripotency exit

We then asked whether subdivision of the basolateral domain into basal and lateral domains, mediated by dynamic interplay between cell-matrix and cell-cell contacts, could play a role in pluripotency exit. We first tested the effect of nocodazole on pluripotency gene expression. Nocodazole can destabilize microtubule network and lead to basement membrane degradation of chicken epiblast cells during gastrulation EMT at stage HH3/4 (17, 40), potentially erasing the difference between the basal and lateral membrane domains. However, nocodazole was also shown to reduce pluripotency marker expression through an upregulation of TP53 in ESCs (41). Treatment of stage HH1 embryo with nocodazole (3 hours; 10 μg/ml) led to disruption of epithelial integrity and to a modest decrease in NANOG protein expression levels (Fig.S4E,F). HiPSCs treated with nocodazole (10 μg/ml) for 24 hours resulted in their detachment from the dish. When treated for 3 hours (10 μg/ml), it caused a mild, but statistically significant drop in NANOG and POU5F1 expression levels (Fig.4L; Fig.4M, red: nocodazole, blue: control. N=3). NANOG and POU5F1 intensities showed tight correlation in both control and nocodazole treated groups (Fig.4N), except for a small population of mitotically arrested cells (Fig.4O, right) which had a strongly reduced level of POU5F1 expression (Fig.4N,O). Taken together, these data suggested that microtubule destabilization had a complex, but not prominent, effect on hiPSC pluripotency exit.

We next tested whether integrin-mediated cell-matrix interaction played a role in pluripotency exit. Both chicken epiblast and hiPSCs expressed β1 integrin as the major β isoform (Fig.1C; Fig.5A; Sup-Table3). The blocking antibody for β1 integrin (P5D2; Materials and Methods) would block β1 integrin-mediated cell-matrix interaction, but not the E-cadherin mediated cell-cell interaction. 24 hour-treatment with the blocking antibody (1 μg/ml) led to complete detachment of hiPSCs, suggesting that it had a potent effect on cell-matrix interactions. 2 hour-treatment of β1 integrin blocking antibody (1 μg/ml) did not cause obvious morphological abnormality, but resulted in a robust upregulation of POU5F1 expression in all colonies and erased the center (-) territory in large colonies (Fig.5B), suggesting that integrin mediated cell-matrix interaction was a key regulator of pluripotency exit. Prolonged treatment (48 hours) of hiPSCs with much lower concentrations of β1 integrin blocking antibody (10 ng/ml and 24 ng/ml), however, resulted in reduction in pluripotency marker expression (Fig.5C). Interestingly, both concentrations (10 ng/ml and 24 ng/ml) of blocking antibody treatment also led to an increase in colony sizes [both center (+) and center (-) types] (Fig.5D), indicating a “delay” in the timing of the appearance of patterning. Conversely, treatment with an integrin beta1 activating antibody (P4G11; Materials and Methods) reduced the colony sizes when patterning started to emerge (Fig.5E). Collectively, these data suggested that integrin-mediated signaling played a positive role in promoting the pluripotency exit. We next asked how E-cad mediated cell-cell interaction was involved in this process. HiPSC colonies treated with EGTA (2mM) for 20 minutes stilled retained overall integrity (Fig.5F; left panels), although cells had rounded-up morphology and longer treatment led to cell detachment. Interestingly, 20 minute EGTA treatment completely abolished patterned expression of pluripotency markers in large colonies, resulting in uniform salt-and-pepper expression of POU5F1 and eliminating the difference between central and pericentral territories (Fig.5G,H; Fig.5I, EGTA top). This loss of patterning was not rescued even after three hours of recovery in normal medium (Fig.5I, EGTA bottom). Because EGTA blocks both E-cad (cell-cell) and Integrin (cell-matrix) signaling, we tested the effect of EGTA treatment in the presence of Mg^2+^ (5 mM) or combined Mg^2+^ (2.5 mM) and Mn^2+^ (2.5 mM). Both Mg^2+^ and Mn^2+^ are known to specifically promote integrin-mediating signaling (42). Addition of Mg^2+^ did not rescue the effect of 20min EGTA treatment (Fig.5G,H; Fig.5I, top), but did rescue the control pattern after 3 hours’ recovery (Fig.5I, bottom). Combination of Mg^2+^ and Mn^2+^ very robustly inhibited the effect of EGTA (Fig.5G,H,I), suggesting that pluripotency exit seen in the center of normal colonies was primarily mediated by through integrin signaling. Further supporting our hiPSC-based observations, freshly-laid avian embryos, in which pluripotency markers were ubiquitously expressed in the epiblast (Fig.5K, control; Fig.S3A and Fig.5L) and pluripotent cells were of partial epithelial morphology (Fig.1), brief activation of the integrin signaling (EGTA+ Mg^2+^+Mg^2+^) also led to dramatic reduction of Nanog expression (experimental outline in Fig.5J; Nanog mRNA in Fig.5K, right; Nanog protein in Fig.5N).

**Figure 5:**
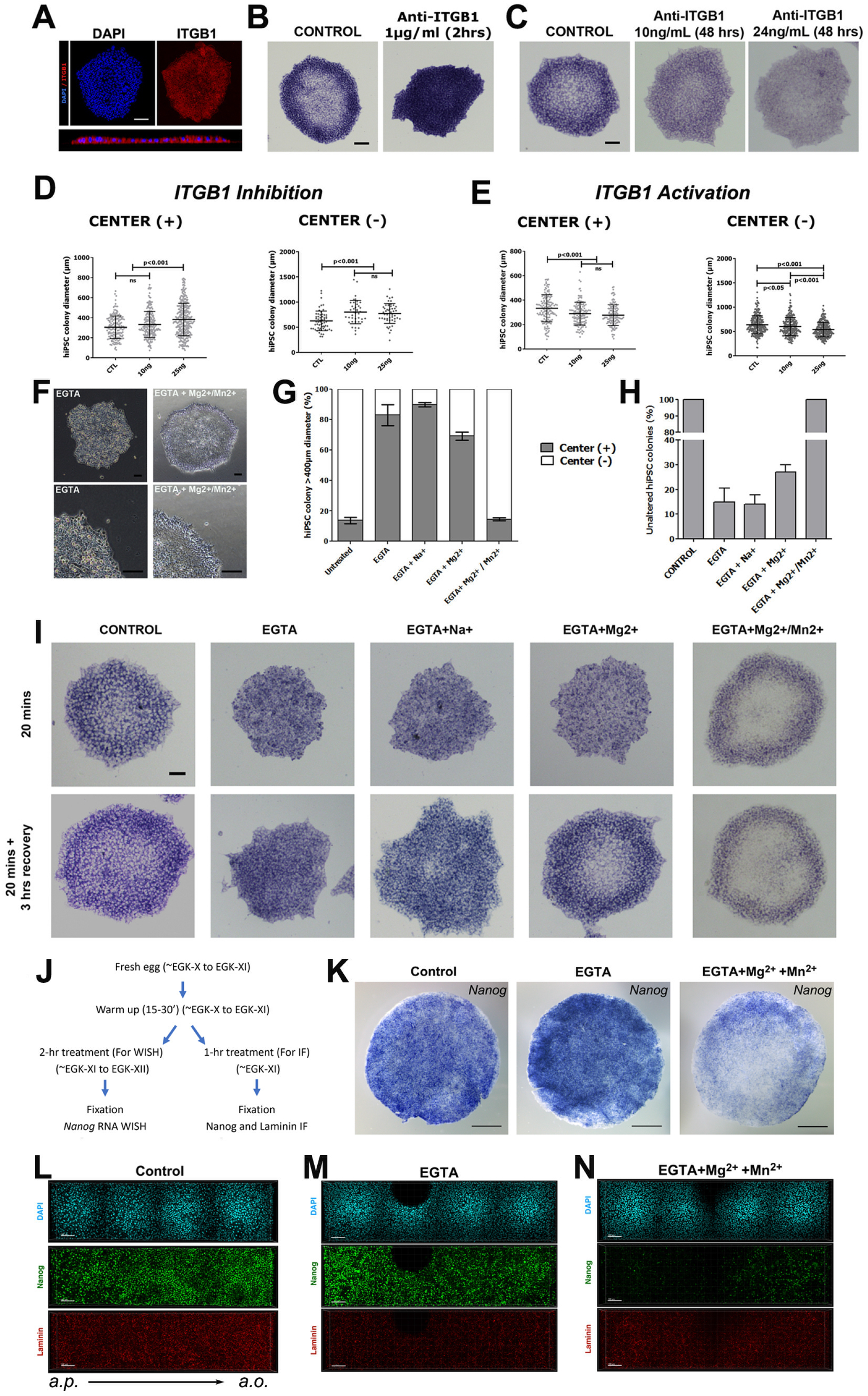
Integrin signaling regulates pluripotency exit. (A) Representative image of Integrin-β1 (ITGB1) expression in large hiPSC colonies, corresponding to the sum of the images of a confocal acquired stack (step 0.75 μm), and related reconstructed median optical cross section of this colony. (B-E) In situ hybridization visualization of POU5F1 mRNA expression under control and ITGB1 perturbation conditions. (B) hiPSC colonies treated with ITGB1 blocking antibody for 2h at 100 ng/ml. POU5F1 RNA in situ. (C) hiPSC were cultured in the presence of 10 ng/ml or 25 ng/ml of ITGB1 blocking antibody for 48h. POU5f1 RNA in situ. (D) Analysis of the mean colony diameter of central (-) and central (+) hiPSC colonies. Addition of 10 or 25 ng/ml of ITGB1 blocking antibody significantly increased the mean size of central (-) colonies (p<0.001) compared to control. 25ng/ml of ITGB1 blocking antibody also increased central (+) ones (p<0.01). (E) Analysis of the mean colony diameter of central (-) and central (+) hiPSC colonies after addition of ITGB1 activating antibody (10 or 25 ng/ml for 48h). Contrary to ITGB1 blocking antibody, ITGB1 activating antibody decreased the mean size of central (-) and central (+) colonies (p<0.001), suggesting that hiPSC epithelialization was initiated in smaller colonies. (F) Bright-field views of hiPSC colonies 20min after treatment either with EGTA alone or with EGTA and Mg2+/Mn2+. Bottom panels: magnified views. (G,H,I) Representative images (I) and quantification (G,H) of POU5F1 mRNA expression in hiPSC colonies after 20 min treatment of either EGTA alone, EGTA and Na+ together, EGTA and MG2+ together, or EGTA and Mg2+/Mn2+ together, compared to the control (left most panel). Top: 20 min treatment; Bottom: 20 min treatment followed by 3h recovery in normal culture medium. Scale bar:100μm. (J) Schematic diagram of treatment of chicken embryos with EGTA plus Mg2+/Mn2+ treatment. Freshly layed eggs were warmed for 15-30 minutes, followed by New culture incubation for 2 hours with albumen replaced by either PBS(+) (control), 2mM EGTA in PBS(+), or 2mM EGTA plus Mg2+/Mn2+ in PBS(+). Embryos were then fixed for whole mount in situ hybridization or immunofluorescence analysis. Both the starting and ending stages were categorized as early HH1. (K) EGTA plus Mg2+/Mn2+ treatment (right) greatly reduced pluripotency marker Nanog expression in area pellucida (central epiblast). EGTA only treatment (center) did not have prominent effect on Nanog expression. Scale bar: 1mm. (L-N) Immunofluorescence analysis of treated embryo with Nanog and Laminin antibodies. Nanog protein expression levels were greatly reduced in 2mM EGTA plus Mg2+/Mn2+ treated embryos (N) whereas EGTA alone did not prominent effect on Nanog protein expression. With either EGTA alone or EGTA plus Mg2+/Mn2+ treatment, the basement membrane (laminin) remembered the control case. Scale bar: 100μm.

### MET-mediated hiPSC pluripotency exit involves canonical EMT/MET transcriptional regulator

Above data suggested that epiblast pluripotency regulation is tightly coordinated with epiblast MET regulation. As in any EMT/MET process, epiblast MET is presumed to be under stereotypic transcriptional regulation. Among the core EMT/MET regulators (SNAI1, SNAI2, ZEB1, ZEB2, TWIST1 and TWIST2) (43), data from geo-profiles suggested that SNAI1, ZEB1 and ZEB2 were highly expressed in hiPSCs (44). We confirmed high expression of SNAI1 in hiPSCs by RNA in situ hybridization analysis (Fig.6A). Interestingly, SNAI1 expression also showed robust patterning in large colonies (Fig.6A, middle), similar to that of pluripotency markers. SNAI2 expression was very low or negative in hiPSCs (Fig.6A, right). This colony-size dependent patterning of SNAI1 expression was also seen at the protein-level, with pan-colony expression in smaller colonies (Fig.6B, top) and patterned (center-low and periphery-high) expression in larger ones (Fig.6B, bottom). Aside from the core EMT/MET transcription factors, additional transcriptional regulators are known to play a role in EMT/MET in a process-specific manner and we tested three (GRHL2, YAP and TAZ) among those. GRHL2 was shown to recruit polycomb repressor complex and suppress ZEB1/2 and other EMT-related genes (45–47) and its down-regulation was associated with poor prognosis in ovarian cancer patients (45, 48). Immunofluorescence staining showed that GRHL2 was highly expressed in hiPSCs, but without exhibiting marked regional difference (Fig.6C). We furthermore tested YAP and TAZ, which are Hippo pathway effectors (49) and are known to be involved in EMT regulation (43). Nuclear localization of YAP was shown to positively control epiblast pluripotency in both human and mouse models (50, 51). Immunofluorescence staining showed that YAP was expressed and localized to the nucleus in all hiPSCs (Fig.6D), with a slight decrease in center cells. TAZ was shown to differentially affect primed PSC’s self-renewal and differentiation via its subcellular localization (52). Consistent with reported role of nuclear TAZ in promoting EMT, immunofluorescence staining with hiPSCs revealed prominent difference in TAZ subcellular distribution, with pericenter cells having a nuclear localization and center cells having a cytoplasmic one (Fig.6D,E) (53). Both SNAI1 and TAZ were speculated to promote cancer EMT through their regulation of matrix metalloproteinases (MMPs), especially MMP9 (54, 55). We investigated MMP2 and MMP9 activities (both are expressed in hiPSCs based on micro-array data) in hiPSC colonies using sequence specific MMP fluorescent sensors (56, 57) (Materials and Methods). MMP9 activity was high throughout hiPSC colony (Fig.6G), whereas MMP2 activity was relatively weak but detectable (Fig.6G). As a negative control, MMP13 activity was not detectable in hiPSCs (Fig.6G). However, neither MMP9 nor MMP2 exhibited prominent patterning in their activity distribution, suggesting that MMP activity is not the primary target of epiblast MET regulation. This was supported by the observation that treatment with Marimastat, a pan-MMP inhibitor, did not lead to significant alteration in POU5F1 patterning in hiPSCs (Fig.6F). Taken together, these data suggested that several EMT/MET transcriptional regulators are involved in the epiblast MET process. How epiblast MET is transcriptionally regulated remains to be clarified.

**Figure 6:**
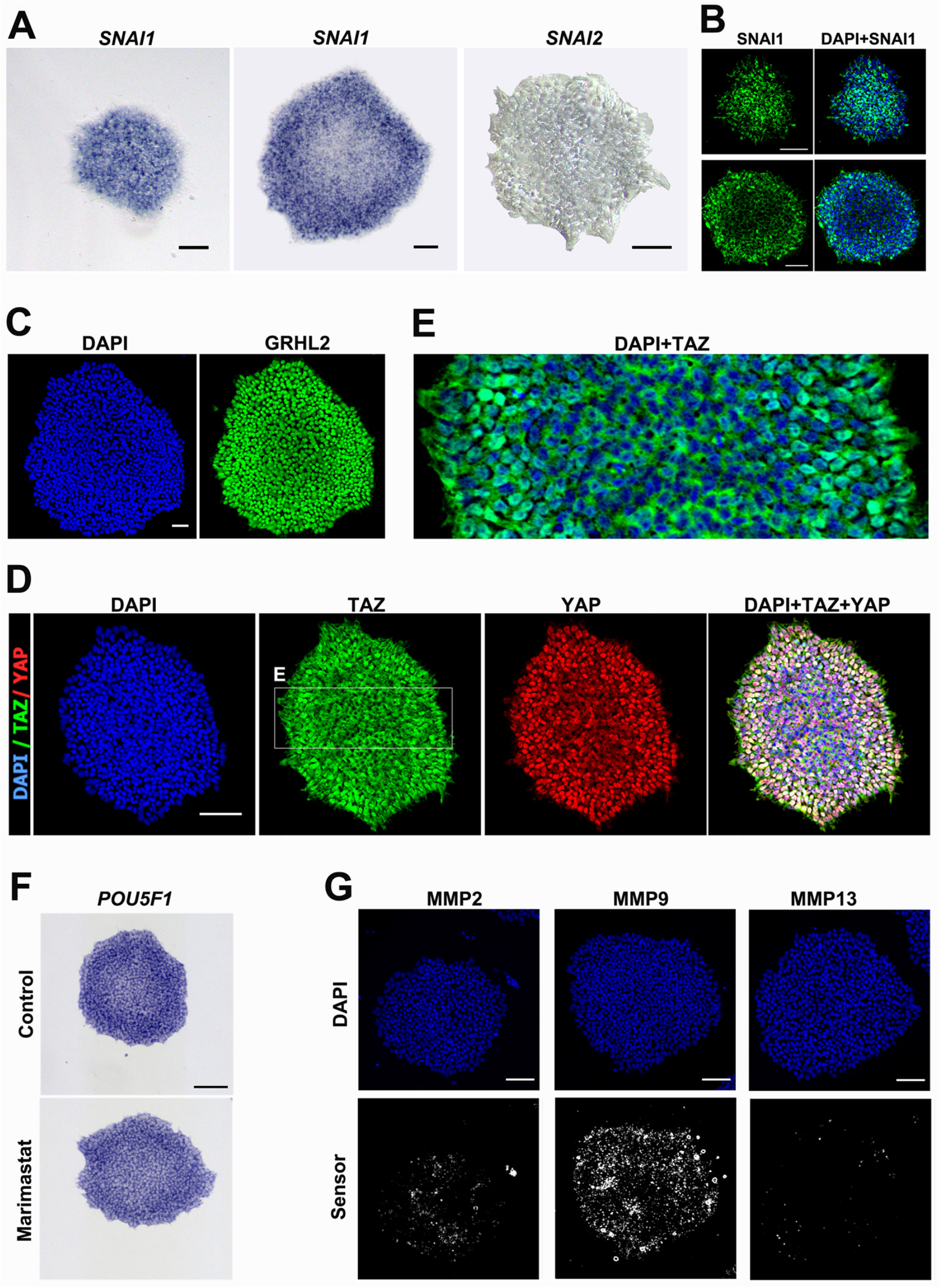
Expression of EMT-related regulators and MMPs in hiPSC colonies. (A) RNA in situ hybridization analysis of SNAI1 (Snail) and SNAI2 (slug) genes. SNAI2 (right) is absent or weakly expressed in hiPSC colonies. SNAI1 (left and middle) is robustly expressed in hiPSC colonies. SNAI1 expression is also patterned (with center low and pericenter high) in large colonies (middle). (B) Immunofluorescence staining of SNAI1, showing uniform expression in small hiPSC colonies and pattern expression (center low and pericenter high) in large colonies. (C) Immunofluorescence staining of GRHL2, showing uniform expression in hiPSC colonies. (D) Immunofluorescence staining of YAP and TAZ, showing uniform expression and subcellular localization for YAP and a clear shift in subcellular localization for TAZ (cytoplasmic in colony center and nuclear in colony pericenter). (E) Magnified view of TAZ ((green) and nuclei (cyan), showing the difference in its subcellular localization between colony center and pericenter regions. (F) POU5F1 RNA expression is not affected by pan-MMP inhibitor Marimastat. (G) Activities of MMPs detected in hiPSCs using specific MMP sensors for MMP2, MMP9 and MMP13 (see also materials and methods). MMP9 is highly active, MMP2 is weakly active and MMP13 is not active. However, intra-colony epithelialization pattern is not reflected in MMP2 activity distribution pattern.

### MET-like colony patterning is also observed in human ESCs, but not in mouse PSCs

We next investigated whether MET-like colony patterning could be observed in other PSCs. Human ESCs (hESCs) were reported to exhibit prepatterning when grown for 24h in micropatterned culture condition (58), with colony size-dependent center-low/periphery-high intra-colony variation in pluripotency marker (NANOG, OCT4 and SOX2) expression. This observation was confirmed when hESCs were cultured in pluripotency maintaining, micropatterned culture condition for either shorter (11h; MP-media11h) or longer (45h; MP-media45h) time period (Fig.7A, top; Fig.7B), with more prominent pre-patterning (periphery high, center low) of NANOG, POU5F1 (OCT4) than SOX2 seen in 45h culture (Fig.7B). Such pre-patterning of pluripotency markers (NANOG and POU5F1) was also observed in hESCs cultured in non-micropattern condition (Dish-media45h; Fig.7A, bottom and Fig.7B). These data suggested that intra-colony pre-patterning is a general behavior of human PSCs. Supporting this, hESC colonies also exhibited a prepatterning of EMT/MET marker SNAI1 (Fig.7A,B), as seen in hiPSC colonies (Fig.6A,B). This MET-like prepatterning of hESC colonies under maintenance conditions is likely different from EMT-like behavior of hESCs induced under BMP-induced differentiation (58, 59). Under BMP-induced differentiation, SNAI1 was highly expressed in colony periphery, and pluripotency markers NANOG and SOX2 were markedly reduced in this region (MP-BMP45h; Fig.S6). Because SOX2 marks both the pluripotent cell population and the neuroectoderm cell lineage, a differentiating hESC colony would show an upregulation, instead of downregulation, of SOX2 in colony center. Under pluripotency maintenance conditions, reduced NANOG and OCT4 expression in colony center was prominent in both hESCs (Fig.7A,B) and hiPSCs (Fig.7C,D,E) and this was coupled with a mind decrease in SOX2 (Fig.7) rather than differentiation-associated increase (Fig.S6), suggesting that pluripotency exit is not correlated with spontaneous differentiation.

**Figure 7:**
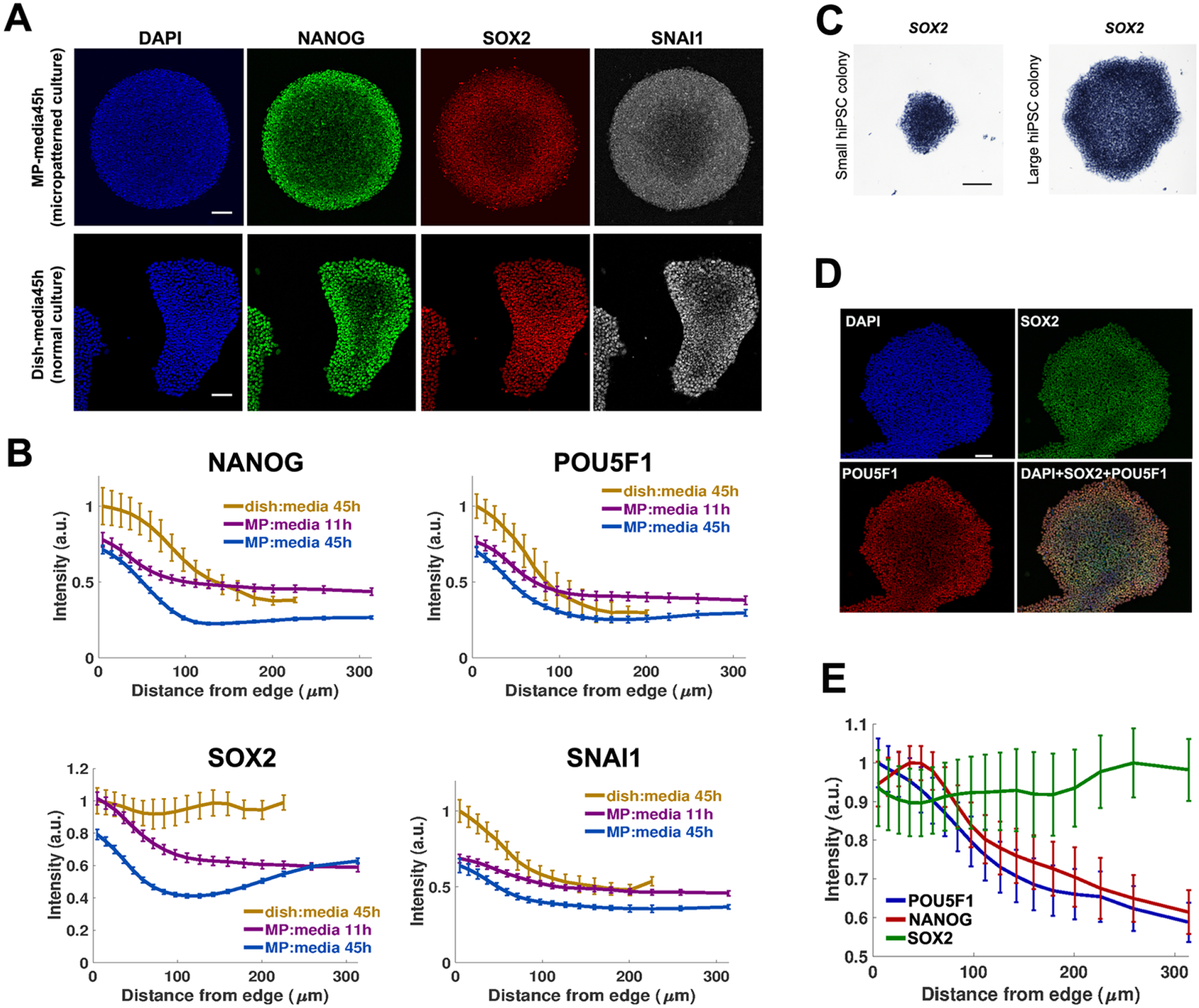
hESCs exhibit similar intra-colony patterning of pluripotency and EMT markers. (A) hESCs were seeded and cultured for 45h in either micro-patterning (MP-media45h) or normal (Dish-media45h) conditions (see Materials and Methods for details). Colonies were then stained for NANOG, SOX2 and SNAI1 expression. Similar to hiPSCs, a pattern of center-low and periphery-high expression could be observed for all three markers. (B) Statistical analysis of data shown in (A). Also included in the analysis were additional marker (POU5F1) and culture condition (11h in micro-patterning conditions [MP-media11h]. Under non-differentiation culture conditions (Dish-media45h, MP-media45h and MP-media11h), a center-low/periphery-high pattern of expression could be seen for NANOG, POU5F1, SOX2 and SNAI1.(C) *SOX2* mRNA expression in small (left) and large (right) colonies. (D) SOX2 protein expression in large colonies (green), co-stained for POU5F1 (red). (E) Statistical analysis of data shown in (D). Also included in the graph was additional marker NANOG. Both NANOG and POU5F1 have prominent pericenter-high and center-low pattern. In either hiPSCs or hESCs, SOX2 does not show up-regulation in colony center, suggesting that these cells are no differentiating into neuroectoderm lineage (as seen with center cells under induced differentiation; see Sup-Fig.6).

The hiPSCs have a transcriptome profile similar to post-implantation epiblast and EpiSCs (60). In vivo, epithelialization of the epiblast corresponds to developmental progression from pre-implantation to post-implantation epiblast(61). We then asked whether similar prepatterning could be observed in mouse EpiSCs (mEpiSCs). We first checked epithelial status of mEpiSC and mESC colonies (Fig.S5A,B). mEpiSC colony (Fig.S5B) had a single cell-layered structure with well-localized ZO-1 (tight junction) and E-cad (adherens junction) with no obvious regional variation, indicating its full epithelial organization. In contrast, mESC colony had a multilayered structure with irregular E-cad and ZO-1 signals (Fig.S5A). mESCs had higher levels of expression of pluripotency markers NANOG and POU5F1 at both the protein (Fig.S5C) and RNA (Fig.S5D) levels. But no intra-colony pre-patterning could be observed in either mEpiSC or mESC colonies. Similar to hiPSCs, GRHL2 was expressed in all cells in both mEpiSC and mESC colonies, but without any overt patterning (Fig.S5E). Interestingly, we did observe a prominent difference in SNAI1 subcellular localization, with predominantly cytoplasmic localization in mESCs and predominantly nuclear localization in mEpiSCs. Taken together, these data suggested that in mouse PSCs under the current culture conditions, differential epithelialization status and pluripotency level was not reflected as intra-colony prepatterning, but rather as differences in colony morphology and pluripotency regulation between mESCs and mEpiSCs.

## DISCUSSION

The avian embryo before gastrulation contains three cell lineages: the epiblast, hypoblast and area opaca (Fig.8A, left). The first two are equivalent to their mammalian counterparts (Fig.8A, right). The third anchors the developing embryo to the yolk and stretches the epiblast as it expands over the yolk surface. The area opaca cells are also the evolutionary origin of mammalian trophoblasts. The monotremes, the earliest branched-out extant mammals, for example, still retain an avian-like pre-gastrula organization (8). Despite superficial difference in epiblast topography between the chick (exposed to the embryo exterior) and human (covered by polar trophectoderm), epiblast cells in both species undergo a very similar sequence of molecular and morphogenetic events (Fig.8A). These events include: 1) molecular specification of epiblast precursors; 2) morphological sorting of epiblast-fated cells; 3) full epithelialization of the epiblast (subject of this study); 4) morphogenesis of epithelialized epiblast; 5) dissolution of epithelial structure during gastrulation EMT. It is generally considered that pluripotency loss coincides with the last step in this sequence when mesendoderm cells switch off pluripotency markers after gastrulation EMT and the remaining epiblast cells differentiate into ectoderm derivatives and likewise lose pluripotency. In this work, we demonstrated that there is an additional component of pluripotency regulation which is distinct from gastrulation EMT-related pluripotency loss. We termed this process “pluripotency exit” and its earliest step “initiation of pluripotency exit”. We provided evidence that this “initiation of pluripotency exit” is correlated to and regulated by epiblast partial MET (Fig.8A). This partial MET refers to the transition from an immature epithelioid organization to a fully-epithelial one. Using a combination of avian epiblast and mammalian PSC models, our data suggested that partial MET-mediated pluripotency exit is reversible and is evolutionarily conserved. It is primarily regulated through the specification of epiblast’s basal membrane domain and by Integrin-mediated epiblast-extracellular matrix signaling.

**Figure 8:**
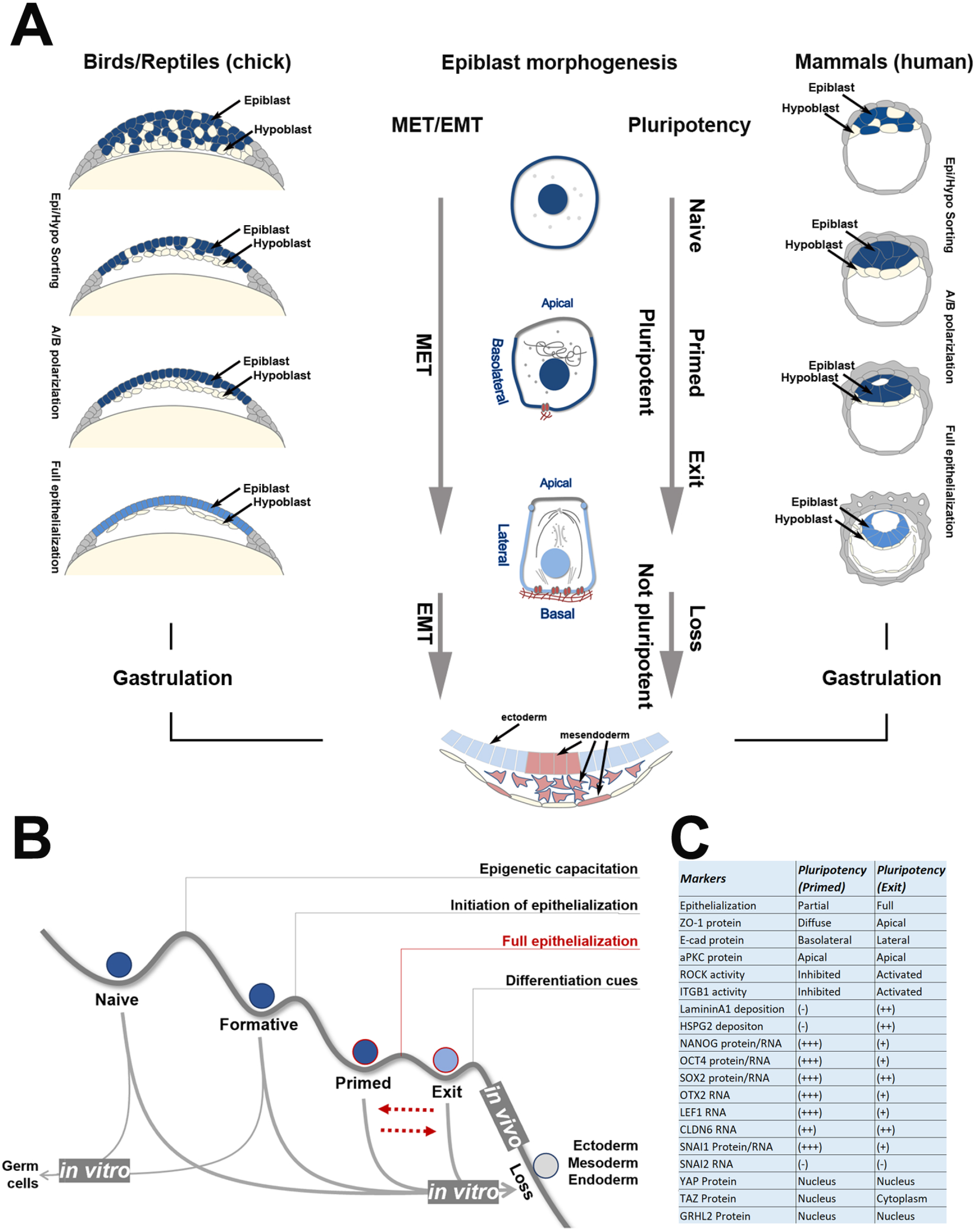
Summary of epiblast MET and pluripotency exit prior to gastrulation-EMT associated pluripotency loss in both human and chicken development. (A) Developmental progression leading to gastrulation in the chick (left) and human (right). Blue: epiblast; light-blue: epithelial epiblast; light-yellow: hypoblast; grey: area opaca/trophectoderm. Epi: epiblast. Hypo: hypoblast. A/B polarization: apicobasal polarization. In amniote development, pre-gastrulation epiblast morphogenesis consists of sequential steps in attainment of epithelial features, from non-polarized epiblast, to partially apicobasally-polarized epiblast, to fully epithelial epiblast. Epiblast cells are considered to be pluripotent in all these MET phases. A precise one to one correlation of in vivo cell biological features and in vitro PSC subtypes may not be possible. But the MET step described in this work likely corresponds to the later phase of primed pluripotency status during which a transition from partially epithelialized epiblast to fully epithelialized epiblast takes place, marked by a reduction in pluripotency (e.g., NANOG and POU5F1) and mesenchymal (e.g., SNAI1) regulators. Such epithelization and pluripotency reduction are reversible and epiblast cells at this phase are still considered pluripotent. Gastrulation, coupled with a full EMT, marks the onset of irreversible pluripotency loss. (B) Comparison of the pluripotency exit state described in the work with other known ones, including the naïve, the formative and the primed states. In vivo, it is hypothesized that pluripotent cells transit through those states sequentially. In vitro, however, each state is capable of direct differentiation into three germ layers. The naïve and formative states are also capable of differentiating into the germ cell fate. (C) A summary of molecular and cellular features that define the pluripotency exit state.

The process of epiblast pluripotency exit described in this work is likely different from the “naïve-to-primed” transition that has been reported in the mouse (62) (Fig.8B). The basic distinctions between these two in vitro (naïve and primed) states of pluripotency are their in vitro culture conditions and in vivo reconstitution potentials. The naïve state has been hypothesized to represent the pre-implantation epiblast, whereas the primed state the post-implantation one (3). Cell biological studies suggest that this transition is associated with polarization of the mouse and human epiblast (62). However, our study showed that in both chicken epiblast at EGK-X (oviposition stage) and young hiPSC colonies, the pluripotent stem cells are already polarized in their apical-basolateral axis, suggesting that the starting point of our analysis is a primed-like state of pluripotency. This hypothesis is supported by molecular data from hiPSCs (showing a primed pluripotency signature) and from our comparative analysis of pluripotency markers in two avian species, the zebra finch (oviposition at stage EGK-Vl; with a prominent naïve pluripotency signature) and chick (oviposition at stage EGK-X; with a primed pluripotency signature) (21). Despite such distinctions, the “naïve-to-primed” transition and the “pluripotency exit” reported here likely represent the same sequence of events in epiblast MET (Fig.8A,B), with the former corresponding to the first half of MET (from non-polarized epiblast to polarized epiblast) and the latter to the second half (from polarized epithelioid epiblast to fully epithelial epiblast). More importantly, it is worth noting that epiblast MET, including both components discussed above, is different from the pluripotency loss seen during gastrulation EMT, and that these two morphogenetic events are separated by a wide (in developmental terms) temporal window during which epiblast cells are known to undergo additional intra- and inter-cellular morphogenesis while keeping their epithelial status. One of the most important patterning events in early amniote development, the radial-symmetry breaking which determines the future embryonic dorsal-ventral axis, takes place during this interval period. It remains to be demonstrated whether there are additional molecular checkpoints between the MET and EMT processes discussed above. A possible scenario is that in vivo, as has been proposed for pluripotent cells cultured in vitro (63) and supported molecularly by the presence of many microstates during ESC differentiation into neuronal lineage (64), a continuum of regulatory steps will be uncovered.

A key component of this MET-associated pluripotency exit is the integrin-mediated cell-matrix signaling. Such signaling requires the deposition of basement membrane proteins, as shown in this work for both the chicken epiblast and hiPSC, and the segregation of basal and lateral membrane domains of the pluripotent cells. We have previously shown that during chicken gastrulation EMT, epiblast cells interact with their basement membrane through Integrins and Dystroglycan (18, 40). This interaction compartmentalizes the basolateral domain of an epiblast cell into separate basal and lateral domains and restricts functions of cell-cell interaction molecules (e.g., E-cad) to the lateral domain and cell-matrix interaction molecules (e.g., Integrins and Dystroglycan) to the basal one. We confirmed that Dystroglycan was also expressed in pre-gastrulation epiblast cells (Fig.1D) and in hiPSCs, although in this work we did not focus our study on the function of Dystroglycan in pluripotency exit. Among the Integrins, in both gastrulation stage epiblast (40) and pre-gastrulation stage epiblast (22) (Sup-Table3), ITGB1, ITGB5 and ITGB3 are the major beta subunits and ITGAV and ITGA6 are the major alpha subunits. Interestingly, Integrin profiling of 201B7 line (Sub-Table3) suggests that such heterogeneity is conserved in hiPSCs, which express ITGB1, ITGB5, ITGAV, ITGA6 and ITGAE as their major alpha and beta subunits, respectively. Involvement of Integrin signaling in pluripotency cell maintenance has been reported. For example, Integrin-mediated focal adhesion kinase activity was shown to protect hESCs from apoptosis and differentiation (65) and alphavbeta5 Integrin was shown to support self-renewal of hESCs (66). Furthermore, alpha6beta1 was shown to mediate laminin-511 supported renewal of hESCs and hiPSCs (67), but likely through alpha6-mediated suppression of beta1 Integrin activity, as activation of beta1 Integrin or knockdown of apha6 Integrin led to a reduction of pluripotency marker expression (68). Supporting this, inhibition of Integrin signaling by disintegrin could promote pluripotency maintenance of mouse iPSCs (69). In our work, brief suppression of Integrin signaling using high concentrations of beta1 Integrin blocking antibody resulted in a dramatic increase in pluripotency level and in the blockage of pluripotency exit (Fig.5B) and brief activation of Integrin signaling using Mg2+ and Mn2+ cations led to pluripotency exit in both hiPSCs (Fig.5F) and chicken embryos (Fig.5K,N), strongly suggesting that MET-associated pluripotency exit observed in our model systems is positively regulated by Integrin signaling. However, our results also pointed to a putatively positive role of Integrin signaling in pluripotency maintenance, because long-term treatment of low concentrations of beta1 Integrin blocking antibody resulted in a reduction of pluripotency marker expression (Fig.5C). Future experiments on how Integrin signaling is transmitted intracellularly to regulate the pluripotency network will be needed to clarify such discrepancy.

Our work is the first to associate epiblast cell morphology with epiblast pluripotency exit in embryonic epiblast and cultured PSCs. It is unclear whether this phenomenon can be generalized to all cultured pluripotent cells that adopt an epithelial-like colony structure during their maintenance phase or whether it reflects intrinsic morphogenetic behavior of their developmental origin, the epiblast. We did not observe intra-colony MET in mEpiSCs, although these cells adopt a perfect epithelial morphology throughout the colony. Such species-specific difference may be due to difference in culture conditions, but it also suggests that morphological feature is not a sole indicator of variability in the pluripotency spectrum. Patterned heterogeneity in pluripotency marker expression was proposed to facilitate higher-order patterning under differentiation conditions in human ESCs and mouse epiblast-like cells (58, 59, 70), possibly by facilitating lineage specific gene expression through GRHL2-mediated enhancing switching in the regulation of pluripotency genes (71). As MET-like pre-patterning was also observed in hESC colonies, it would be interesting to test in the future whether difference in differentiation potentials can also be associated with cells isolated from sub-regions of hESC colony with different epithelialization status and EMT/MET marker expression levels. Such a hypothesis is supported by a recent paper which showed that intra-colony difference in cellular adhesive properties defines a potential N-cadherin+ founder cell population within primate PSC colonies (72).

More importantly, as we have demonstrated, pluripotency exit is reversible and is different from pluripotency loss seen during gastrulation in vivo or under differentiation conditions in vitro. For ectoderm-fated epiblast cells, the MET is clearly necessary because both neural ectoderm and non-neural ectoderm cells adopt a fully epithelial morphology after gastrulation. For those epiblast cells which will eventually give rise to the mesoderm and endoderm germ layers during gastrulation, they will have to undergo an MET to achieve full epithelial status only to be followed by a full EMT and loss of their pluripotency. The biological significance for such morphogenetic behavior is unclear because in vitro, lineage differentiation into the three germ layers can be achieved from pluripotent cells regardless of their epithelialization status (Fig.8B). Although the term epiblast generally refers to pluripotent cells in amniotic vertebrates (birds, reptiles and mammals), pluripotency regulatory network has deep evolutionary root (73) and similar pluripotent cells also exist in the anamniotic vertebrates [e.g., in zebrafish (74, 75)]. It is therefore interesting to note that full epithelialization of pre-gastrulation pluripotent cells only occurs within the amniote clade (8, 76). This suggests that epiblast MET is associated with certain amniote-specific developmental features which require epithelial structure-based intercellular signaling or force transmission [e.g., epiblast cell intercalation and polonaise movement reported in the pre-gastrulation chick embryo (77–79)]. These epithelial structure-based cellular events are known to be directly involved in the formation of the primitive streak (an amniote-specific feature) or more generally, of a posterior epiblast-restricted center of mesendoderm internalization (i.e., of later-on gastrulation EMT). Therefore, epiblast MET is not only important for epiblast pluripotency regulation as described in this work, but likely also for epiblast planar symmetry-breaking. How these two phenomena are linked developmentally remains to be clarified.

## MATERIALS & METHODS

### Array analysis of chicken area pellucida samples

Fertilized chicken eggs purchased from a local farm (Shiroyama farm, Kanagawa, Japan) were incubated without storage at 38.5°C for 0.5, 5.5, 9 and 14 hours. The area pellucida (AP) region of the embryos, containing both the epiblast and hypoblast, but excluding the primitive streak region for 9 and 14 hour sample sets, was cut out in Pannett-Compton solution. Dissected AP samples of specific time points were pooled and used for RNA isolation. Two independent sets of total RNA from pooled AP samples were isolated using QIAGEN RNeasy Micro Kit. 100 ng of quality checked total RNAs were used for array analysis, with the cDNA synthesis and cRNA labeling reactions performed according to the two-cycle protocol provided by Affymetrix. Affymetrix high-density oligonucleotide arrays for Gallus gallus (GeneChip^®^ Chicken Genome) were hybridized, stained, and washed according to the Expression Analysis Technical Manual (Affymetrix) and analysis of resulting expression values was performed as described previously (80). Array raw data have been deposited in public database with the accession number GSE114476.

### hiPSC culture and reagents

The hiPSC line 201B7 (purchase from RIKEN BRC Cell Bank)(was used for all experiments. Cells were maintained in an undifferentiated state on iMatrix-511 (0.5 μg/cm^2^; Nippi, Japan; #892018)-coated dish using the StemFit AK02N (Ajinomoto) medium and cultured at 37 °C with 5% CO_2_. At Day 7 of culture, hiPSC colonies were dissociated using accutase (Nacalai tesque, Japan; #12679-54) and unicellular hiPSCs were then seeded on new culture dish freshly coated with iMatrix-511, in a medium containing 10 μM Y-27632 (Wako Pure Chemicals Industries, Japan; #253-00513). All experiments in this study were performed at day 6 of culture, one day before normal passaging. For pattern alteration experiments, Y-27632 was used at a concentration of 20 μM, MMP broad spectrum inhibitor Marimastat (a gift from Prof. Hiroshi Sato, Kanazawa University) at 150 nM, nocodazole (Wako #140-08531) at 10 μg/ml, EGTA (Sigma #E3889) at 2 mM, MgCl2 (Sigma #1901905) at 5 mM when alone or at 2.5 mM when associated to MnCl_2_ (2.5mM; Wako #139-00722), and NaCl at 5 mM (Wako #195-01663). Monoclonal antibody against β1-Integrin (inhibiting antibody clone P5D2 or activating antibody clone P4G11; Developmental Studies Hybridoma Bank #AB528308 or #AB528307) was added to the culture media either at day 4 of culture at the concentration of 10 ng/ml and 25 ng/ml or for only 2h prior to fixation at 1 μg/ml. These antibodies were also added to PBS (-) supplemented with 2.5 mM of MgCl_2_.

### hESC culture

ES017 hESCs were grown in mTeSR1 (STEMCELL Technologies) in tissue culture dishes coated with Matrigel (Corning; 1:200 in DMEMF12) overnight at 4°C. Cells were passaged using dispase (STEMCELL Technologies) every 3 days. Cells were routinely tested for mycoplasma contamination and found negative. Micropatterning experiments were also performed with mTeSR1 following the protocol previously described (2). Where indicated, cells were treated with 50 ng/ml BMP4 to induce differentiation.

### mPSC culture

The mESC line EB5 was grown on 0.1% gelatin-coated dishes in a medium consisting of GMEM 10% knock-out serum replacement, 1%FBS, 1 mM sodium pyruvate (Wako #190-14881), 1x non-essential amino acids (Sigma #M7145) and LIF 1000u/ml. Cells were passaged every three days using accutase. The mEpiSC line (81) was grown on 0.5 μg/cm^2^ iMatrix-511 (Nippi, Tokyo, Japan) coated dishes in a medium consisting of DMEM-F12, 20% knock-out serum replacement, 1 mM sodium pyruvate, 1x non-essential amino acids, 10^-4^ M 2-ME, 10 ng/mL activin A (R&D Systems #338-AC), 5 ng/mL human recombinant FGF2 (R&D Systems #3139-FB) and 10μM of XAV939 (Abcam #ab120897). Cells were passaged every 4-5 days using accutase for single cell dissociation.

### Laser microdissection

hiPSC colonies were grown on plasma treated glass bottom dish coated with iMatrix-511. At day 6 periphery and center of patterned colonies were separated using the laser beam of a Zeiss PALM microbeam. Following laser microdissection, both regions were isolated manually under a microscope. Selected cells were treated with Y-27632 for 2h prior to dissociation and re-plated either at a density of 200 cells/cm^2^ for pluripotency assessment experiments or at 400 cells/well (Prime Surface 96V Sumilon Sumitomo Bakelite, Wako #MS-9096V) for differentiation experiments.

### hiPSC differentiation and PCR analysis

After five days of culture, newly formed embryoid bodies differentiation was induced by replacing maintenance media with DMEM 10% FBS. After three days, samples were collected for RNA extraction (RNeasy micro Kit, QIAGEN) along with MCF-7, HepG2, HUVEC and undifferentiated hiPSC that were used as positive and negative controls. PCR was performed by using the following primers, human GAPDH: 5’-TCATCCCTGAGCTGAACGGG-3’ (forward) and 5’-TCCCCTCTTCAAGGGGTCTACA-3’ (reverse); human NESTIN: 5’-CAGGGTTGGAACAGAGGTTGG-3’ (forward) and 5’-GCATCTACAGCAGGAGAGGGTG-3’ (reverse); human SOX1: 5’-CGGAGCTCGTCGCATTTGTT-3’ (forward) and 5’-TCCCCGGGGTTCCCTTACTT-3’ (reverse); human PAX6: 5’-CACCCGCCCTGGTTGGTATC-3’ (forward) and 5‘-TGAGGGCTGTGTCTGTTCGG-3’ (reverse); human FOXA2: 5’-AGCGGTGAAGATGGAAGGG-3’ (forward) and 5’-ATGGCCATGGTGATGAGCGA-3’ (reverse); human T: 5’-CCGAGAGCGCGGGAAAGAG-3’ (forward) and 5’-TCACTATGTGGATTCGAGGCTCAT-3’ (reverse); human HAND1: 5’-TTAACAGCGCATTCGCGGAG-3’ (forward) and 5’-CGTGCGATCCAAGTGTGTGG-3’ (reverse); human EOMES: 5’-ACACTTTACCTCAAGCCCGC-3’ (forward) and 5’-AGTTGCTAGGAGACAGCCGC-3’ (reverse); human GATA4: 5-‘GTCCCAGTGCAGACCTGCTG-3’ (forward) and 5’-CCCTGAGGCTGTAGGTTGTGT-3’ (reverse); human CDX2: 5-‘CTTCCTGCGCTTCTGGGCT-3’ (forward) and 5’-CCAGGCACTGAGGCTTGC-3’ (reverse).

### In Situ-Hybridization

Probes used for In Situ-Hybridization (ISH) were obtained by RT-PCR using the following primers for human NANOG: 5’-GTGTGGATCCAGCTTGTCC-3’ (forward) and 5’-GTCACACCATTGCTATTCTTC-3’ (reverse) 493bp; human POU5F1: 5’ -CAAGAACATGTGTAAGCTGCGG-3’ (forward) and 5’-AGGAGTACAGTGCAGTGAAGTG-3’ (reverse) 425bp; human LEF1: 5’-CCAGACAAGCACAAACCTCTC-3’ (forward) and 5’-AGCCAAGAGGTGGGGTGATC-3’ (reverse) 420bp; human OTX2: 5’-GAGAGGACGACGTTCACTC-3’ (forward) and 5’-TCTGACAGTGGGGAGATGG-3’ (reverse) 365bp, human SNAI1: 5’-TGCCTCGACCACTATGCCG-3’ (forward) and 5’-AGGCTCGAAAGGCCTTCAACT-3’ (reverse) 479bp; human SNAI2 5’-TTCGTAAAGGAGCCGGGTGAC-3’ (forward) and 5’-ATCTTTGGGGCGAGTGAGTCC-3’(reverse) 425bp; human CLDN6: 5’-ACTCGGCCTAGGAATTTCCCTT-3’ (forward) and 5’-TAATCCCCGTGTGCTGGACG-3’ (reverse) 473bp; human MMP2: 5’-TCTTTGGACTGCCCCAGACA-3’ (forward) and 5’-AGTACTCCCCATCGGCGTTC-3’ (reverse) 447bp; human MMP9 5’-AAGGCCAATCCTACTCCGCC-3’ (forward) and 5’-AGGGCGAGGACCATAGAGGT-3’ (reverse) 445bp; mouse POU5F1 5’-CAGGACATGAAAGCCCTGCAGAA-3’ (forward) and 5’-GCCCAAGCTGATTGGCGAT-3’ (reverse) 397bp; and mouse NANOG 5’-CTGGGAACGCCTCATCAATGC-3’ (forward) and 5’-TACTCCACTGGTGCTGAGCC-3’ (reverse) 469bp. In situ-hybridization was performed as described previously for chick embryos (82) with slight adaptation in incubation time or concentration of reagents. More specifically, cultured cells were fixed in 4% paraformaldehyde overnight at 4°C. Permeabilization step was performed using 1 μg/ml proteinase K incubation for 20 minutes at room temperature, followed by post-fixation for 15 minutes, prehybridization at 68°C for 3 hours, and hybridization with gene specific antisense DIG labelled probes at 68°C overnight. After hybridization, samples were washed in prehybridization solution at 68°C and then in TBST at Room temperature. This was followed by 1 hour of blocking at RT, overnight incubation in anti-DIG antibody solution (1/2000; Roche #11093274910) at 4°C. Finally, after several washing steps in NTMT solution, hiPSC colonies were processed for color development in NBT and BCIP at room temperature shielded from light. Whole mount in situ hybridization with chicken embryos for Nanog mRNA detection was carried out as previously decribed (82) and probe information for chicken Nanog was reported previously (19).

### Imaging and Immunofluorescence staining for PSCs

All immunofluorescence experiments were performed using cells grown on plasma treated glass bottom dish. The following primary antibodies were used for NANOG (1:500; ReproCELL #RCAB004P for hiPSCs, 1:200 R&D Systems AF1997 for hESCs or #RCAB001P for mouse PSCs), OCT3/4 (aka POU5F1; 1:500; BD Transduction Laboratories #611202), SOX2 (1:200; Cell Signaling #3579 for hESCs and 1:1000 with rabbit anti sera against mouse Sox2 #164), Omni-probe D8 (1:200; Santa Cruz Biotechnology #sc-7270), E-cadherin (1:500; BD Transduction Laboratories #610181), ZO-1 (1:100; Invitrogen #40-2200), β-Dystroglycan (1:100; Leica #NCL-b-DG), aPKCς clone C-20 (1:400; Santa Cruz Biotechnology #sc-216), β1-Integrin clone P5D2 (1:100; Developmental Studies Hybridoma Bank #AB528308). LAMA1(1:200 SIGMA #L9393), HSPG2 (1:200 Milipore #MABT12), GRHL2 (1:500 Sigma #HPA004820), SNAI1 (1:200 R&D Systems #AF3639), YAP (1:100 Santa Cruz Biotechnology #sc-101199) and TAZ (1:100 Atlas Antibodies #HPA007415) (both YAP and TAZ antibodies were kindly provided by Dr. K. Nishiyama of Kumamoto University). The following secondary antibodies were used for multicolor detection: Alexa Fluor 488, 568 (1:850; Invitrogen) except for hESCs where the secondary antibodies were used at a concentration of 1:500 (Alexa Fluor 488, 555 and 647). Images were acquired using an Olympus FV1200-IX-KU laser scanning confocal microscope for all samples except for 3D reconstruction where a Leica TCS SP8 confocal microscope was used. Single cell 3D reconstruction experiments were generated using the Imaris software v8.0 (Bitplane), and all other image analyses were performed using Fiji software.

### Imaging and Immunofluorescence staining for chicken embryos

Chicken embryos were fixed in 4% PFA. For whole mount staining of Nanog, embryos were processed for immunofluorescence staining, then cleared with SeeDB 2G tissue-clearing solution (83) before being processed for imaging. The following primary antibodies were used, E-cadherin (1:100; BD Transduction Laboratories #610181), ZO-1 (1:100; Thermo Fisher #40-2200), aPKC clone C-20 (1:100; Santa Cruz Biotechnology #sc-216), GM130 clone 35 (1:100; BD Bioscience #610822), AcTub clone 6-11B-1 (1:1000; SIGMA #T6793), β1-Integrin (1:300; Chemicon #MAB13443), Dystroglycan clone H242 (1:100; Santa Cruz biotechnology #sc-28535), Pan-Laminin-1 clone 3H11 (1:100; DSHB #AB528342), Laminin-A1 (1:100; SIGMA #L9393), Agrin clone 6D2 (1:100; from DSHB), NANOG (1:500; kindly provided by Dr. Agata from Gakushuin University, Japan) (84). The following secondary antibodies were used for multicolor detection: Alexa Fluor 488, 568 or 594 (1:300 for whole mount embryo staining, 1:500 for frozen section staining; Invitrogen). Images were acquired using Olympus FV1000 with BX61WI upright or FV3000RS with IX83 invert laser scanning confocal microscope with UPlan-SApochromat 60X/1.2 NA, UPLSAPO 60X/1.3 NA or UPLSAPO 30XS/1.05 NA objective lenses.

### HiPSC sample processing and histology

HiPSC line 201B7 was cultured in the same condition as described above, but on Poly carbonate membrane inserts (Nunc #140660). At Day 6, membranes were washed with PBS and fixed in 4% paraformaldehyde for 15 minutes, followed by a second washing step using PBS membrane and coating with 1% bovine Gelatin (Sigma #G9391) for 1 hour at 37°C. Membranes were then washed and fixed again overnight in 4% paraformaldehyde. Counterstaining of hiPSC colonies was done using H&E staining reagents. Briefly, membranes were incubated with Meyer-Hematoxyline (Sakura Finetek Japan #8650) for 5 minutes, and Eosin (Sakura Finetek, Japan; #8659) for 2 minutes. Membranes were then dehydrated using methanol and isopropanol serial baths before paraffin permeabilization (3x 40 minutes each). Membranes were then cut away from the inserts using a scalpel, and placed in a mold for final paraffin-embedded sectioning (10 μm). Sections were collected on bovine Gelatin coated slides, dewaxed and mounted. Analysis was performed using an Olympus BX51 microscope equipped with a DP70 digital camera.

### Expression constructs and transfection

The pEF-His-A-derived mammalian expression vector, T7/His epitope-tagged human wild type PAR1b/MARK2, “kinase deficient” PAR1b-KM (K49M) and non-phosphorylatable PAR1b-TA (T595A) constructs were kindly provided by Dr. Shigeo Ohno (Yokohama City University, Japan) (39). All transfections have been performed using the Gene Juice^®^ reagent (EMD Millipore #70967) following manufacturer instruction at day 4 of hiPSC culture.

### Synthesis and characterization of MMP-2, −9, and −13 peptide sensors

The MMP sensors consist of MMPs peptide substrate, fluorescence dye (Cyanine 5.5,Ex/Em 675/690 nm), and dark quencher (BHQ-3, abs 650 nm). The MMP-2, −9, or −13 sensor was prepared as previous described (56, 57). Briefly, Cy5.5 succinimide ester and BHQ-3 succinimide ester was conjugated to the MMP-2 [Gly-Pro-Leu-*Gly-Val*-Arg-Gly-Lys-Gly-Gly], MMP-9 [Gly-Lys-Gly-Pro-Arg-*Ser-Leu*-Ser-Gly-Lys-Gly-Gly], and MMP-13 [Gly-Val-Pro-Leu-*Ser-Leu*-Thr-Met-Gly-Lys-Gly-Gly] substrate, respectively, and purified by reverse-phase high-performance liquid chromatography (RP-HPLC). The sensors specificity was confirmed by activated recombinant enzymes (MMP-1, −2, −3, −7, −9, −10, and −13) and fluorescence signal was detected using fluorometer.

### Statistical Analysis

Results were expressed as mean ± SEM. Statistical significance was determined using student’s t test, ANOVA with Bofferoni post test, or Pearson correlation wherever appropriate. Results were considered significant when p<0.05.

## ACKNOWLEDGEMENTS

We would like to thank Ms. Erike Sukowati, Dr. Yuping Wu and Ms. Ayako Isomura for help with chicken embryology, Dr. Hiroki R Ueda, Ms. Junko Nishio and Mr. Kenichiro Uno for help with genechip data production, Dr. Shigeo Ohno, Dr. Tomonori Hirose and Dr. Shoukichi Takahama for sharing PAR1b expression constructs, Dr. Ruby Huang for sharing GRHL2 antibody, Dr. Kiyokazu Agata for sharing chicken NANOG antibody, Dr. Koichi Nishiyama for sharing the YAP and TAZ antibodies, Dr. Hiroshi Sato for sharing the MMP inhibitor Marimastat, Ms. Yangye Zhang and Mr. Galym Ismagulov for help with integrin and EMT marker analysis and Ms. Marina Lizio for help with CAGE data analysis. We would also like to thank the core imaging facility at International Research Center for Medical Sciences, Kumamoto University for help with image acquisition. This work was partially supported by the core research grants to GS from RIKEN Center for Developmental Biology and from International Research Center for Medical Sciences, Kumamoto University; a post-doctoral fellowship to SH from International Research Center for Medical Sciences, Kumamoto University; and by a Takeda research foundation grant to GS.

## SUPPLEMENTARY FIGURE AND TABLE LEGENDS

**Figure S1:**
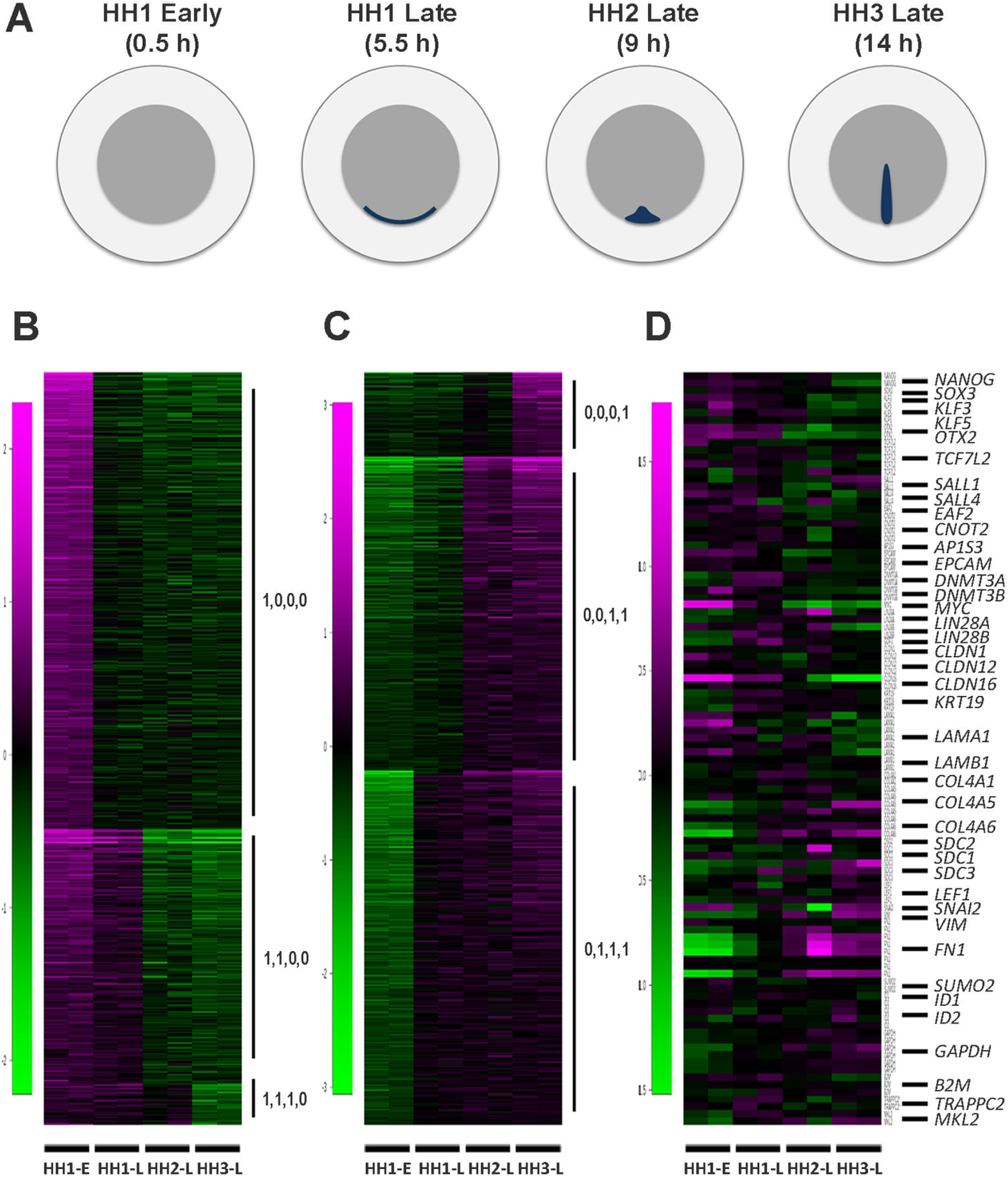
Transcriptomic analysis of pre-gastrulation chicken epiblast. (A) Schematic representation of epiblast tissues used for the analysis. HH1 early: 0.5 hour of incubation; HH1 late: 5.5 hours; HH2 late: 9 hours; HH3 late: 14 hours. Only dark grey areas were collected for Affymetrix chicken genechip analysis. (B-D) Clustering analysis of cross-stage variations, showing significantly down-regulated genes (B), significantly upregulated genes (C) and genes associated with pluripotency regulation and epithelial morphogenesis (D). Each stage is represented by duplicate samples.

**Figure S2:**
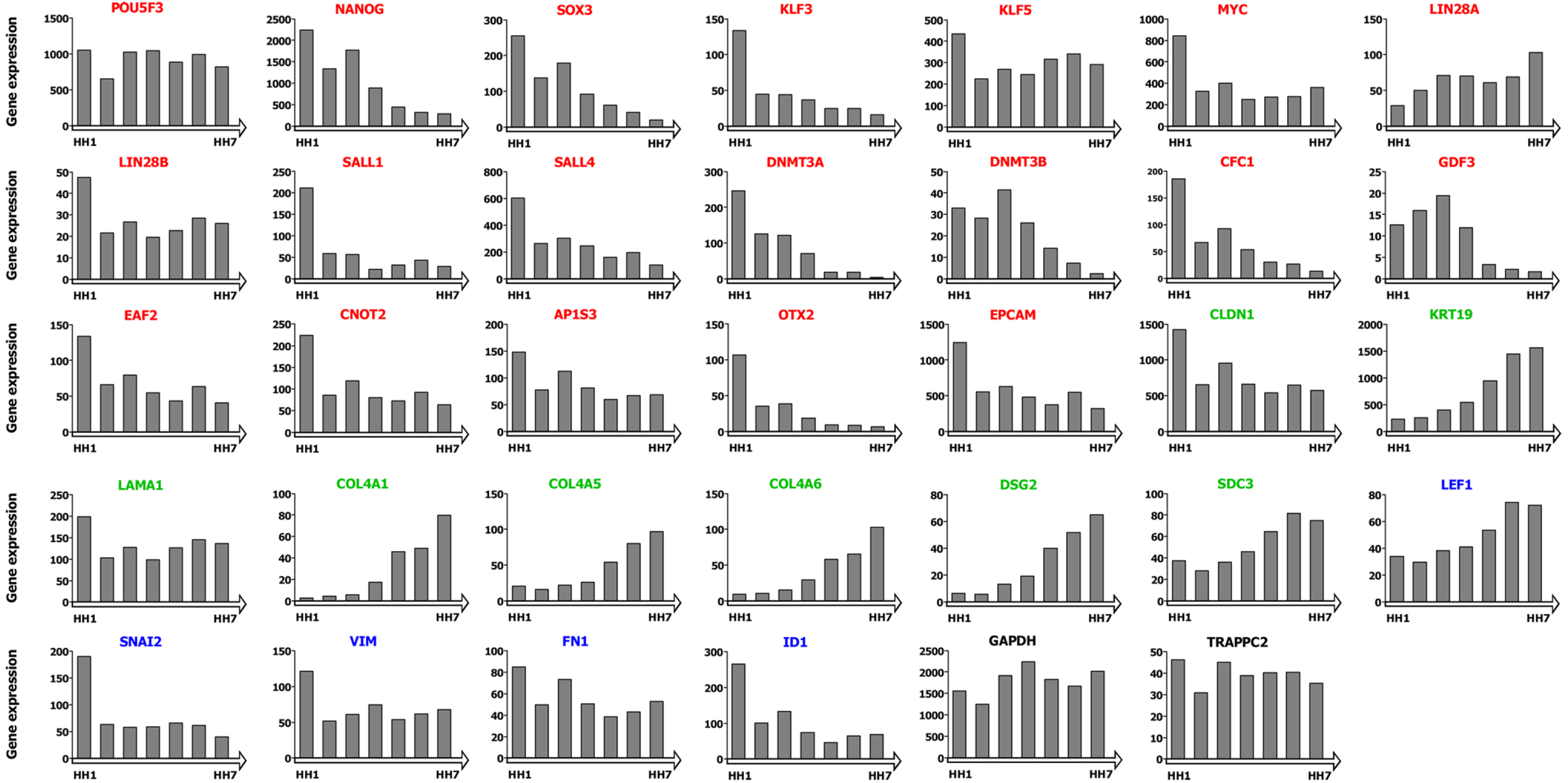
CAGE-based promoter activity of selected genes in early chicken development, from HH1 to HH7. Raw values were extracted from the chicken ZENBU database. Gene names are listed above each panel. Red: pluripotency-related genes; Green: genes related to epithelial features; Blue: genes related to mesenchymal features; Black: housekeeping genes.

**Figure S3:**
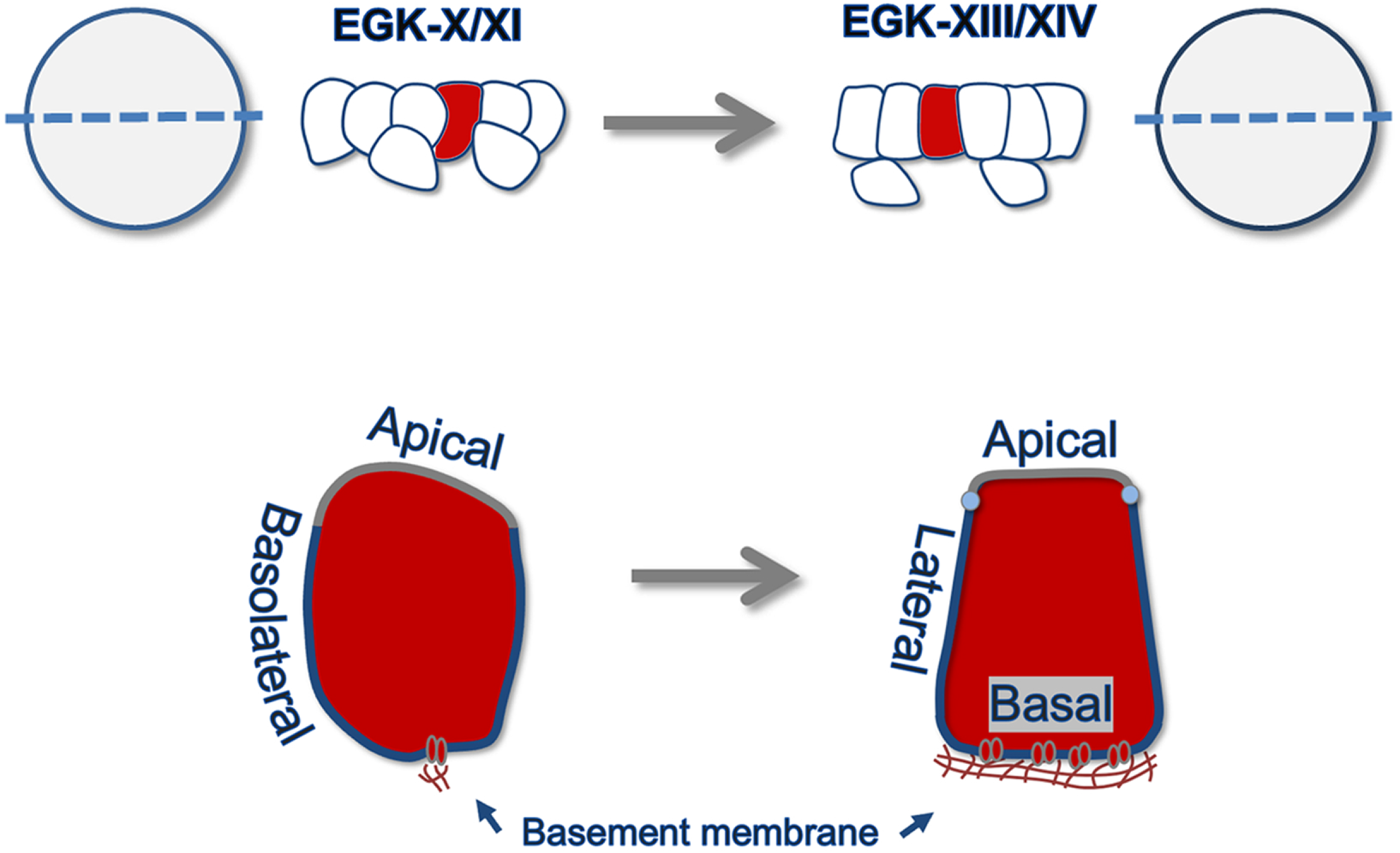
Schematic representation of epiblast morphology at EGK-X/XI and EGK-XIII/XIV. Epiblast cells at EGK-X/XI are partially polarized apicobasally. They become fully polarized by the end of stage HH1 (EGK-XIV), with a continuous layer of basement membrane, smooth apical surface, and segregation of basolateral membrane into basal and lateral compartments.

**Figure S4:**
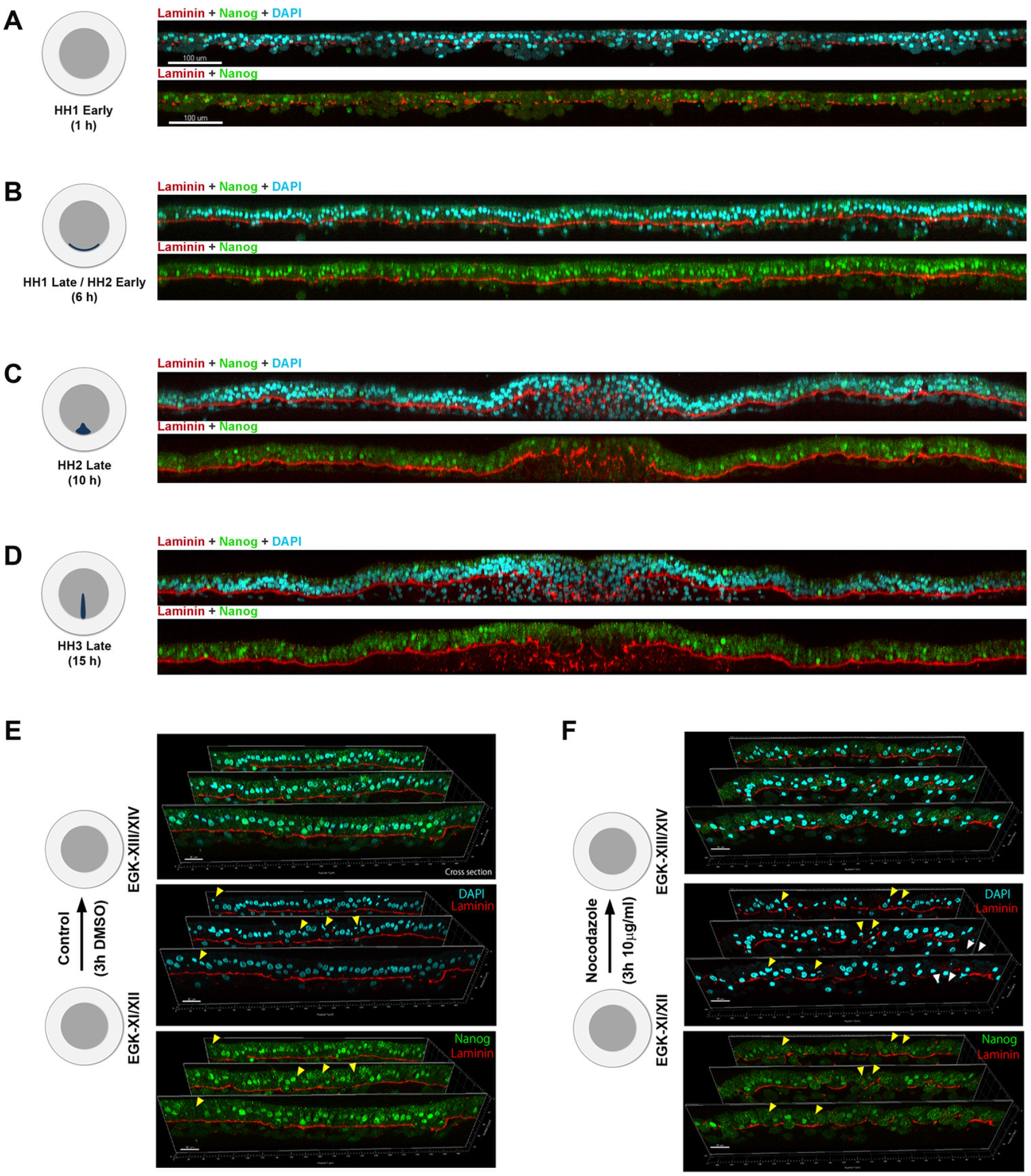
Partial MET during avian epiblast morphogenesis correlates with a progressive pluripotency loss. (A-D) Representative confocal images of embryo sections stained for Laminin (red) and NANOG (green). A: early HH1 (1 hour of incubation); B: late HH1/early HH2 (6 hours); C: late HH2 (10 hours); D: late HH3 (15 hours). In panels C and D, gastrulation EMT is visible in the primitive streak. Scale bar in A-D: 100 μm. (E,F) Control and nocodazole treated embryos stained for NANOG and Laminin. (E): Embryos were pre-incubated for 4 hours under normal conditions and then treated with control (DMSO) for 3 hours. Epiblast cells express NANOG robustly in all control epiblast cells (except in dividing cells, arrowheads). (F): Embryos were pre-incubated for 4 hours under normal conditions and then treated with nocodazole for 3 hours. Nocodazole treatment resulted in Laminin (red) breakdown and in a reduction of NANOG (green) expression level. Nocodazole treatment lowers NANOG nuclear expression (with some examples indicated by arrawheads). Scale bar in E, F: 20 μm.

**Figure S5:**
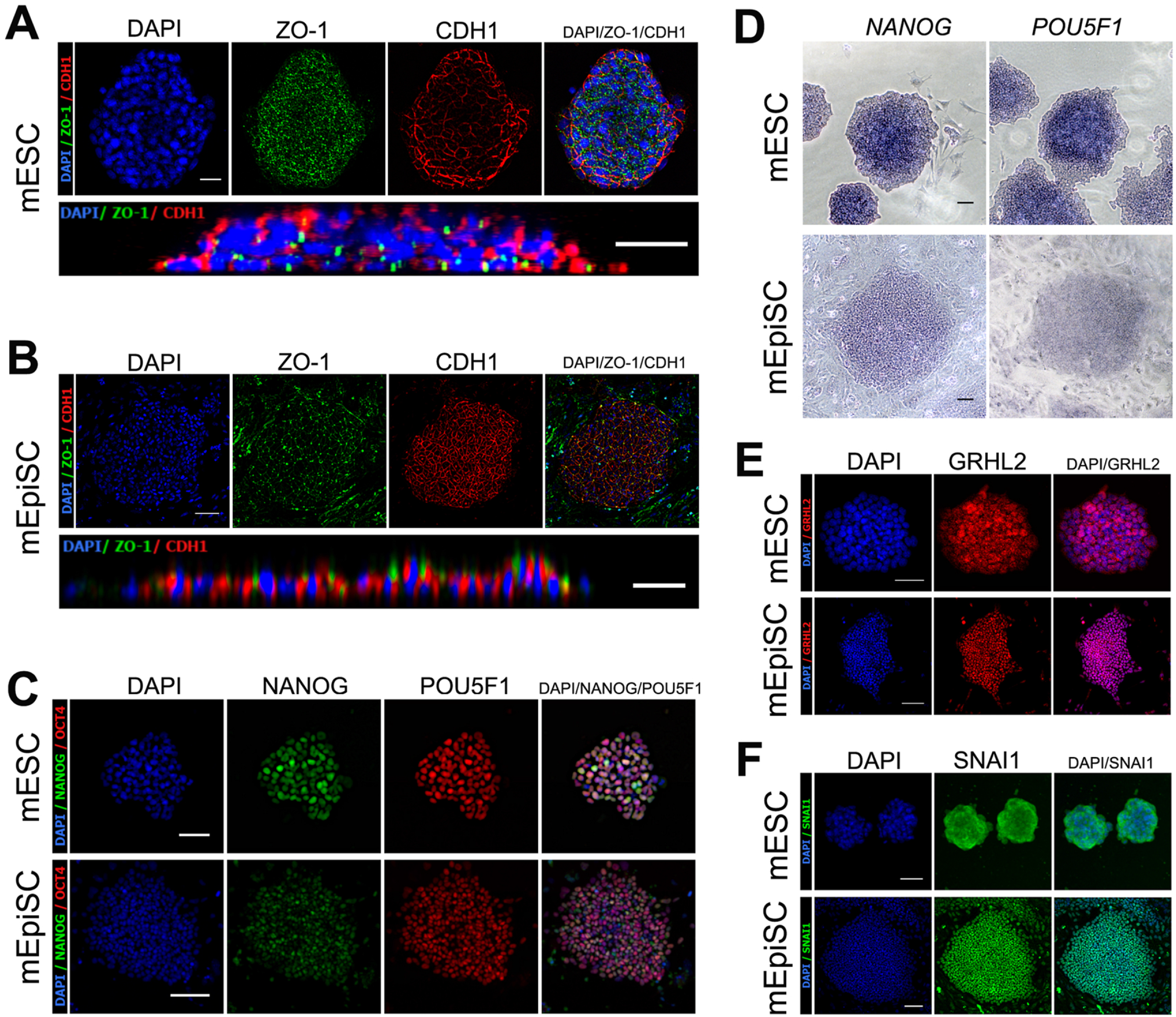
Mouse ESCs and mouse EpiSCs represent two extremes of morphological diversity in PSCs. Mouse EpiSCs (mEpiSCs) were organized as an epithleial structure and expressed lower levels of pluripotency markers. Mouse ESCs (mESCs) were organized as a multilayer non-epithelial structure and expressed higher levels of pluripotency makers. (A) mESC colonies were immuno-stained for ZO-1 and CDH1 (E-cadherin). mESC colonies expressed both ZO-1 and CDH1 and exhibited multilayered organization. Both ZO-1 and CDH1 were poorly localized and mESCs were not organized as an epithelial structure. (B) mEpiSC colonies were immuno-stained for ZO-1 and CDH1 (E-cadherin). mEpiSC colonies exhibited single-cell layered epithelial organization, with proper junctional localization of ZO-1 and CDH1. (C) Pluripotency markers (NANOG and POU5F1) did not exhibit intra-colony patterning in mPSCs. However, protein expression levels of NANOG and POU5F1 were higher in mESCs than in mEpiSCs. (D) RNA in situ hybridization analysis of NANOG and POU5F1 in mESCs and mEpiSCs. RNA expression levels were higher in mESCs than in mEpiSCs. (E, F) Immunofluorescence staining of mESCs and mEpiSCs with GRHL2 and SNAI1 antibodies. Neither GRHL2 (E) nor SNAI1 (F) showed prominent intra-colony patterning. However, SNAI1 was seen to be localized in the cytoplasm in mESCs and in the nucleus in mEpiSCs.

**Figure S6:**
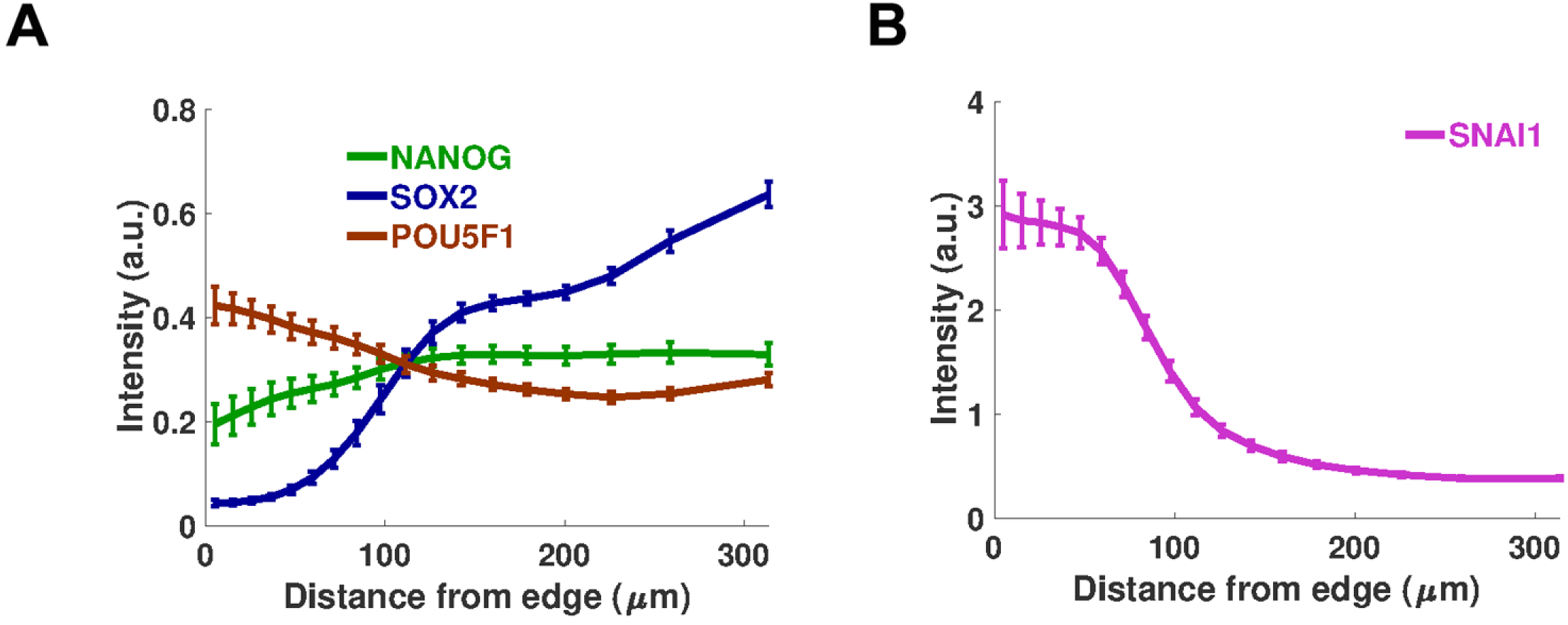
BMP-induced differentiation of micropatterned hESCs. (A) Both NANOG and POU5F1 show strong decrease upon BMP-induced differentiation (compared with expression levels shown in Fig.7B). In contract, SOX2 shows an increase in colony center upon BMP treatment, marking neuroectoderm lineage differentiation. (B) SNAI1 shows dramatic increase in colony pericenter, after BMP treatment, marking mesendoderm differentiation.

**Table S1: Chicken epiblast Affymetrix genechip data.**

**Table S2: Gene ontology analysis of significantly changed epiblast genes from HH1 to HH3.**

**Table S3: Expression levels of Integrin genes obtained from promoterome and transcriptome analyses.** CAGE-based promoter activity data for chicken samples are based on Lizio et al, 2017. Chicken transcriptome data are based on Affymetrix dataset presented in this work. CAGE-based promoter activity data for human iPSCs are based on (85) and human transcriptome data are based on Affymetrix dataset presented in (44).

